# Methylthio-alkane reductases use nitrogenase metalloclusters for carbon-sulfur bond cleavage

**DOI:** 10.1101/2024.10.19.619033

**Authors:** Ana Lago-Maciel, Jéssica C. Soares, Jan Zarzycki, Charles J. Buchanan, Tristan Reif-Trauttmansdorff, Frederik V. Schmidt, Stefano Lometto, Nicole Paczia, Jan M. Schuller, D. Flemming Hansen, Gabriella T. Heller, Simone Prinz, Georg K. A. Hochberg, Antonio J. Pierik, Johannes G. Rebelein

**Affiliations:** Microbial Metalloenzymes Research Group, Max Planck Institute for Terrestrial Microbiology; Marburg, Germany; Department of Chemistry, RPTU Kaiserslautern-Landau; Kaiserslautern, Germany; Department of Biochemistry and Synthetic Metabolism, Max Planck Institute for Terrestrial Microbiology; Marburg, Germany; Evolutionary Biochemistry Group, Max Planck Institute for Terrestrial Microbiology; Marburg, Germany; Department of Structural and Molecular Biology, Division of Biosciences, University College London; London, UK; Department of Chemistry, Philipps–University Marburg; Marburg, Germany; Metabolomics Core Facility, Max Planck Institute for Terrestrial Microbiology; Marburg, Germany; Center for Synthetic Microbiology (SYNMIKRO), Philipps–University Marburg; Marburg, Germany; The Francis Crick Institute; London, UK; Central Electron Microscopy Facility, Max Planck Institute of Biophysics; Frankfurt am Main, Germany

## Abstract

Methylthio-alkane reductases convert methylated sulfur compounds to methanethiol and small hydrocarbons, a process with important environmental and biotechnological implications. These enzymes are classified as nitrogenase-like enzymes, despite lacking the ability to convert dinitrogen to ammonia, raising fundamental questions about the factors controlling their activity and specificity. Here, we present the first molecular structure of the methylthio-alkane reductase, which reveals large metalloclusters, including the P-cluster and the [Fe_8_S_9_C]-cluster, previously only found in nitrogenases. Our findings suggest that distinct metallocluster coordination, surroundings, and substrate channels, determine the activity of these related metalloenzymes. This study provides new insights into nitrogen fixation, sulfur-compound reduction, and hydrocarbon production. We also shed light on the evolutionary history of P-cluster and [Fe_8_S_9_C]-cluster-containing reductases emerging prior to nitrogenases.

## Main Text

Sulfur is an essential element in amino acids, vitamins and protein cofactors. Bacteria assimilate inorganic sulfur through the uptake and reduction of sulfate^1^. However, under sulfate unavailability, bacteria scavenge organic sulfur via methionine-salvage pathways (MSPs) by recycling metabolic by-products to methionine^2,3^. *Rhodospirillum rubrum* utilizes an anaerobic MSP to convert (2-methylthio)ethanol (MT-EtOH) to methanethiol, direct precursor to methionine, via the methylthio-alkane reductase^4,5^. *In vivo*, this enzyme produces methanethiol from several volatile organic sulfur compounds (VOSCs) besides MT-EtOH, including dimethyl sulfide (DMS) and ethyl methyl sulfide (EMS), releasing in a 1:1 stoichiometry ethylene (C_2_H_4_), methane (CH_4_) or ethane (C_2_H_6_), respectively^4,5^. These small hydrocarbons affect our climate but also offer a sustainable alternative to fossil fuel-based biogas and plastic production. Methylthio-alkane reductase is a new oxygen-independent biogenic source of C_2_H_4_^6^, making it an exceptional option for the production of this important chemical building block. Moreover, methylthio-alkane reductases explain the observed accumulation of C_2_H_4_ in waterlogged anoxic soils^7^ and thus are potentially involved in agricultural yield losses due to inhibitory effects of C_2_H_4_ on plant growth^8^.

Methylthio-alkane reductases have been phylogenetically classified as nitrogen fixation-like (Nfl)-enzymes of the nitrogenase superfamily^4,9^ (Extended Data Fig. 1). Nitrogenases are the only enzymes known to reduce the stable triple bond of dinitrogen (N_2_) to produce bioavailable ammonia (NH_3_)^10^ and distinguish themselves from Nfl-enzymes in their catalytic metal centers^11^. Nfl-enzymes such as the dark-operative protochlorophyllide *a* oxidoreductase or the Ni^2+^-sirohydrochlorin *a*,*c*-diamide reductase, involved in carbon-carbon double bond reductions, only contain [Fe_4_S_4_]-clusters and tetrapyrrole binding sites^12^. In contrast, nitrogenases are known to contain large and complex metalloclusters that allow N_2_-fixation, including the P-cluster, an [Fe_8_S_7_]-cluster, and the iron-molybdenum cofactor (FeMoco), a [MoFe_7_S_9_C-(*R*)-homocitrate]-cluster located in the active site of the molybdenum (Mo)-nitrogenase^11,13,14^ (Extended Data Fig. 1). FeMoco is assembled by the maturase Nif(EN)_2_, a nitrogenase family member which converts an Fe-only precursor, the [Fe_8_S_9_C]-cluster (L-cluster), into FeMoco by replacing a terminal Fe atom with Mo and attaching (*R*)-homocitrate^15,16^ (Extended Data Fig. 1). Distinct activities of nitrogenases and Nfl-enzymes are attributed to these differences in metallocluster content and thus there is great interest to identify the metal centers dictating the carbon-sulfur (C-S) bond reduction activity in methylthio-alkane reductases. Here, we determined the molecular structure of the methylthio-alkane reductase complex by anaerobic single-particle cryogenic electron microscopy (cryo-EM) yielding a global resolution of 2.75 Å. We found that the catalytic component contains P-clusters and [Fe_8_S_9_C]-clusters, making it the first Nfl-enzyme with large nitrogenase cofactors, but unable to reduce N_2_. Our findings suggest that the protein scaffold, rather than the metalloclusters alone, dictates the differing catalytic activities of nitrogenases and methylthio-alkane reductases, advancing our understanding of the origin and evolution of N_2_-fixation.

## Results

### Structural characterization of the methylthio-alkane reductase

To reveal the molecular machinery responsible for C-S bond reduction of methylthio-alkane reductases, we heterologously produced subunits MarH, MarD and MarK from *R. rubrum* and its associated maturation enzyme MarB in an engineered *Rhodobacter capsulatus* strain^17,18^, lacking nitrogenase genes. The reductase and catalytic components of the methylthio-alkane reductase were purified separately using affinity chromatography (Fig. 1a). We determined the oligomeric state of each component by mass photometry: the reductase component forms a MarH_2_ homodimer with a measured mass of 69 kDa (Fig. 1d, top panel; theoretical mass: 64 kDa), while the catalytic component forms a Mar(DK)_2_ heterotetramer with a measured mass of 227 kDa (Fig. 1d, bottom panel; theoretical mass: 217 kDa). In conclusion, methylthio-alkane reductase components have the same stoichiometry as the Mo-nitrogenase components.

**Fig. 1:**
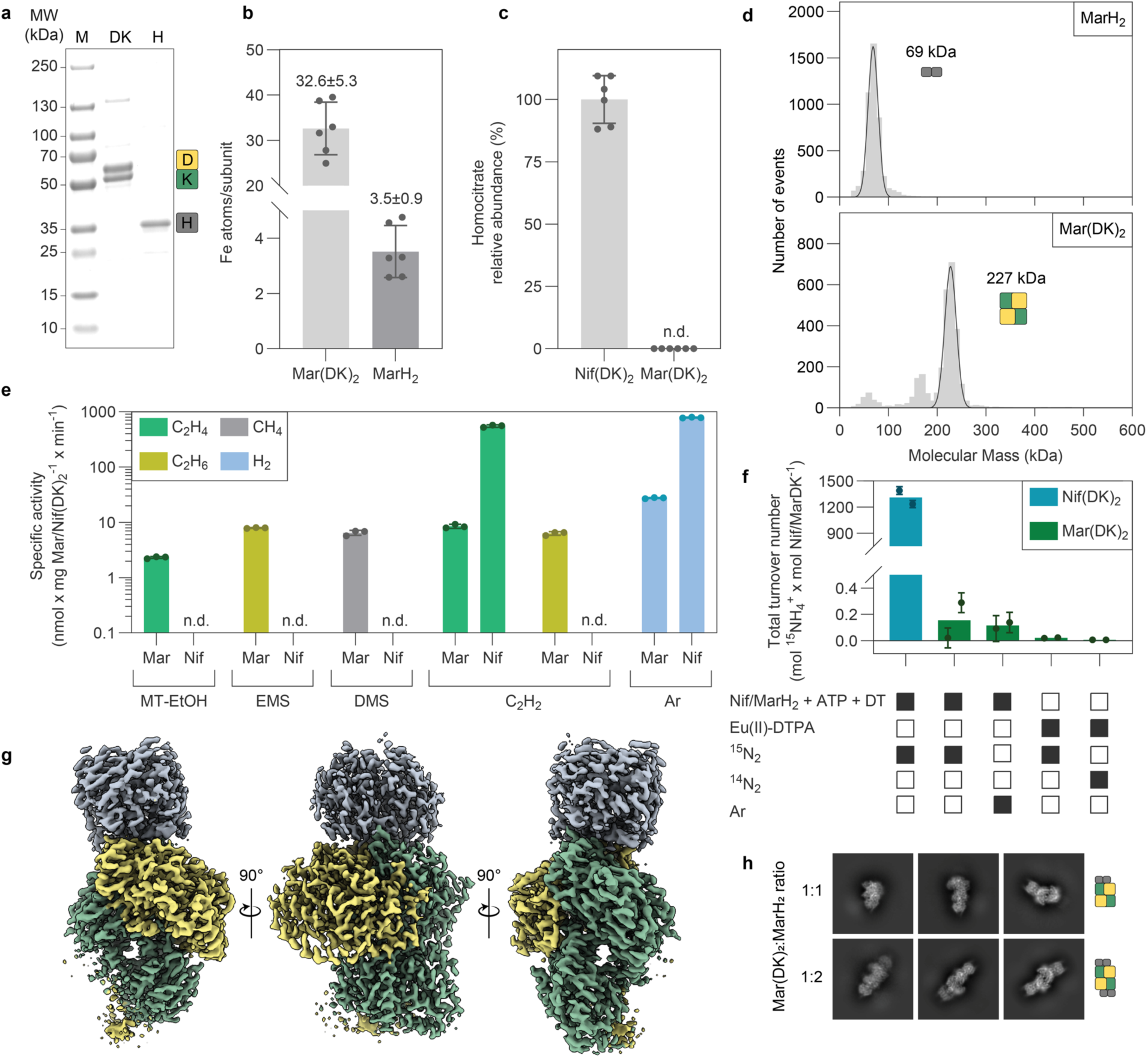
Purification and characterization of the methythio-alkane reductase. **a**, SDS-PAGE of Strep-tagged MarD (57.5 kDa), MarK (50.8 kDa) and His-tagged MarH (31.9 kDa). MW, molecular weight. M, marker. **b**, Fe content of Mar(DK)_2_ and MarH_2_ determined by ICP-OES (n=6). **c**, (*R*)-homocitrate abundance in Nif(DK)_2_ and Mar(DK)_2_ determined by HPLC-MS (n=6). **d**, Mass photometry histogram of MarH_2_ (top panel) and Mar(DK)_2_ (bottom panel). Event counts are plotted against their corresponding molecular mass. **e**, *In vitro* activities of the methylthio-alkane reductase (Mar) and Mo-nitrogenase (Nif) for the reduction of (2-methylthio)ethanol (MT-EtOH), ethyl methyl sulfide (EMS), dimethyl sulfide (DMS), acetylene (C_2_H_2_) and protons under an argon (Ar) atmosphere (n=3). **f**, The methylthio-alkane reductase cannot reduce ^15^N_2_ *in vitro*. Total turnover numbers for the formation of ^15^NH_4_^+^ by Nif(DK)_2_ or Mar(DK)_2_ depending on reductase component, ATP and sodium dithionite (DT), or the strong reductant europium-II-diethylenetriamine pentaacetic acid (Eu(II)-DTPA) detected by NMR, for more details see Extended Data Fig. 7. Columns show the mean, dots the individual measurements (n=2) and error bars the propagated instrumental error. **g**, Electron density map of the MarDK_2_H_2_-complex at a global resolution of 2.75 Å (EMD-50553), contoured at level 10. Reductase component is colored in gray (H_1_ and H_2_), and the catalytic component is colored in yellow (D_1_) and green (K_1_ and K_2_). **h**, Single particle cryogenic electron microscopy (cryo-EM) 2D classes of the methylthio-alkane reductase with ratios of catalytic:reductase component of 1:1 and 1:2. Error bars depict standard deviation unless otherwise stated. n.d., not detected.

The purified enzymes were active for the reduction of MT-EtOH, EMS and DMS as measured by the release of C_2_H_4_, C_2_H_6_ and CH_4_, respectively (Fig. 1e and Extended Data Fig. 2). While the Mo-nitrogenase is unable to reduce VOSCs, the methylthio-alkane reductase converts MT-EtOH at a rate of 2.3 nmol C_2_H_4_ x mg Mar(DK)_2_^-1^ x min^-1^, EMS at 7.9 nmol C_2_H_6_ x mg Mar(DK)_2_^-1^ x min^-1^ and DMS at 6.5 nmol CH_4_ x mg Mar(DK)_2_^-1^ x min^-1^ (Fig. 1e). The methylthio-alkane reductase activity is ATP-dependent and requires the addition of both reductase and catalytic components as well as sodium dithionite (DT) as an electron donor (Extended Data Fig. 2e). Intriguingly, we detected the formation of molecular hydrogen (H_2_) for all tested substrates and in the absence of substrate under an argon atmosphere (Fig. 1e and Extended Data Fig. 2a-d,f). The following reaction stoichiometry was established for MT-EtOH:

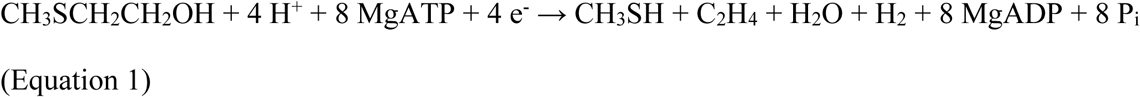

While Mo-nitrogenase cannot reduce VOSCs, methylthio-alkane reductase can reduce the alternative nitrogenase substrate acetylene (C_2_H_2_) to C_2_H_4_ (Fig. 1e and Extended Data Fig. 2d). The permanent formation of H_2_ and the reduction of C_2_H_2_ indicates a highly similar electron transport mechanism in both enzymes. Moreover, methylthio-alkane reductase can fully reduce C_2_H_2_ to C_2_H_6_, a characteristic of the alternative nitrogenases, vanadium (V)- and iron (Fe)- nitrogenases^19,20^, which contain V or Fe in the position of Mo in FeMoco, termed FeVco and FeFeco^21^. This suggests the presence of FeFeco/FeVco-like cofactors in the methylthio-alkane reductase.

To reveal the metalloclusters of Mar(DK)_2_ involved in catalysis, we determined the molecular structure of the methylthio-alkane reductase complex by anaerobic single particle cryo-EM. We were unable to obtain the structure of the catalytic component alone, due to severe preferential orientation bias and susceptibility to denaturation of MarD through contact with the air water interface. We overcame this issue by forming a complex of Mar(DK)_2_ with MarH_2_, using the non-hydrolyzable MgATP mimic MgADP-AlF_3_, thus protecting the catalytic component from denaturation. The complex exists in two major forms: an octameric Mar(DK)_2_(H_2_)_2_-complex, and an hexameric Mar(DK)_2_H_2_-complex (Fig. 1h). By applying strict 3D classification using a local mask on MarD, we achieved an electron density map of the MarDK_2_H_2_ subcomplex with a global resolution of 2.75 Å (Fig. 1g and Extended Data Fig. 3). This allowed us to build a structural model with well-resolved metalloclusters (Fig. 2a), revealing the molecular basis of the methylthio-alkane reductase specificity.

**Fig. 2:**
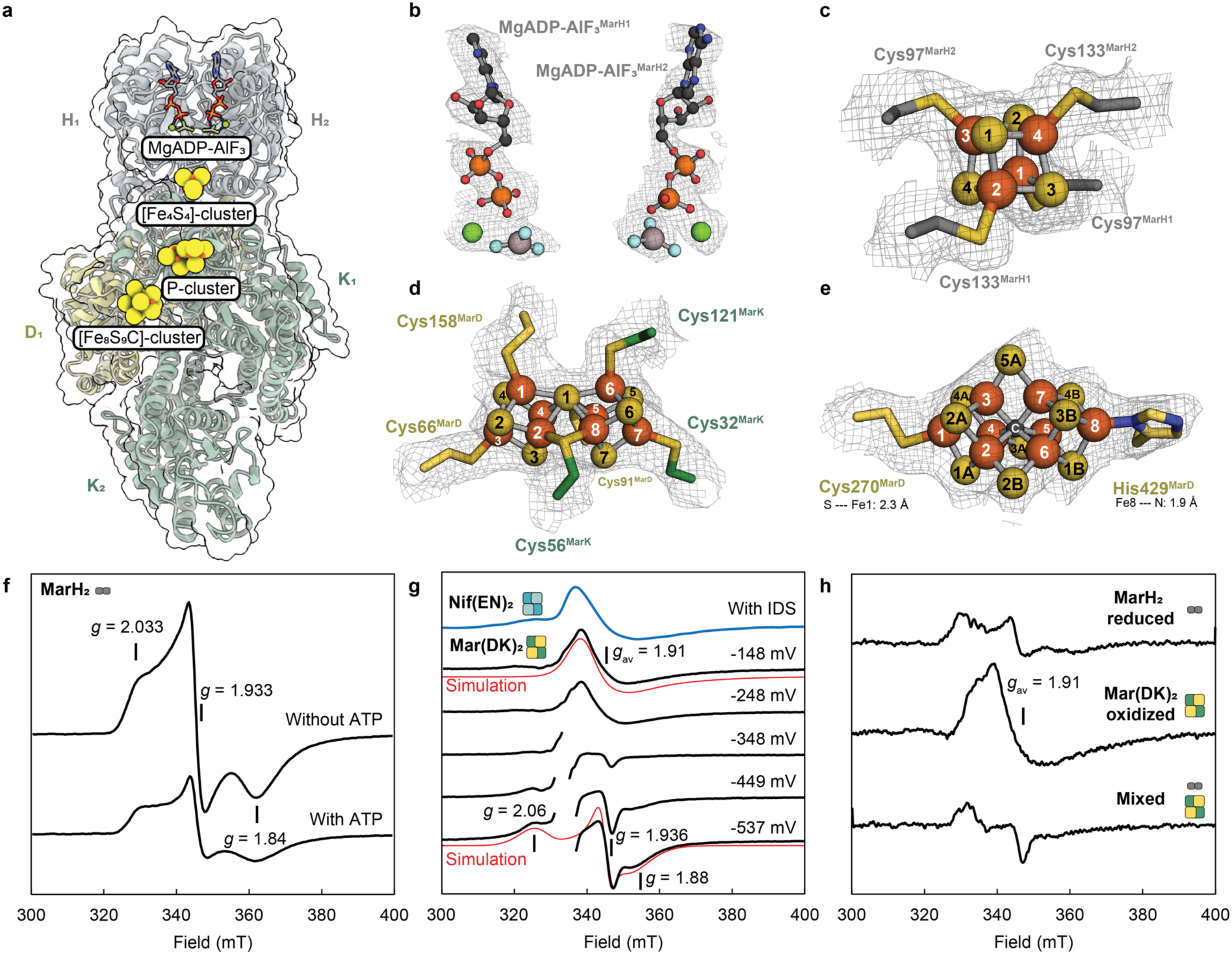
Methylthio-alkane reductase contains nitrogenase cofactors in the active site. **a**, Protein model of the MarDK_2_H_2_-complex with highlighted ligand positions (PDB: 9FMG). Coloring as in Fig. 1g. **b-e**, Magnified views of the methylthio-alkane reductase MgADP-AlF_3_ moieties (b), the [Fe_4_S_4_]-cluster (c), the P-cluster (d) and the [Fe_8_S_9_C]-cluster (e). Cryo-EM electron density maps are displayed as a gray mesh at contour level 8. Ligands are represented as ball-and-sticks with carbon atoms colored in dark gray, nitrogen in blue, oxygen in red, phosphorus in orange, aluminum in light gray, fluoride in pale cyan, magnesium in green, sulfur in yellow and iron in dark orange. Amino acid residues coordinating the metalloclusters are shown in stick representation. **f-h**, EPR spectroscopy of MarH_2_ and Mar(DK)_2_ metalloclusters. (f) EPR *S=*1/2 signals of the [Fe_4_S_4_]^1+^ cluster in dithionite-reduced MarH_2_ with and without MgATP recorded at 10 K. (g) EPR spectra at 10 K of Mar(DK)_2_ samples poised at the indicated redox potentials show two main species between −550 and −150 mV, and with simulations in red lines. The signal appearing upon oxidation is similar to that of the [Fe_8_S_9_C]-cluster of *A. vinelandii* Nif(EN)_2_ (blue trace). (h) EPR spectra of reduced (DT-free) MarH_2_ (upper trace), oxidized (IDS-free) Mar(DK)_2_ (middle trace) and a mixture of both (lower trace) at 10 K. Reduced MarH_2_ can transfer electrons to oxidized Mar(DK)_2_ leading to the disappearance of the broad [Fe_8_S_9_C]-cluster-like *g=*1.91 EPR signal, with concomitant appearance of a sharp rhombic species. EPR conditions: microwave frequency 9.35 GHz; modulation frequency, 100 kHz; 1.5 (f,g) and 1.0 (h) mT modulation amplitude; microwave power 0.2 mW. An isotropic radical was subtracted from all spectra to facilitate simulation and comparison.

### Methythio-alkane reductase contains nitrogenase cofactors

Analyzing the structure, we observed a canonical reductase component MarH_2_, as detailed in the Supplementary Discussion 1, containing one MgADP-AlF_3_ moiety per monomer (Fig. 2b and Extended Data Fig. 4) and a single [Fe_4_S_4_]-cluster in its dimeric interface coordinated by Cys97^MarH^ and Cys133^MarH^ (Fig. 2c). Electrons are transferred from the [Fe_4_S_4_]-cluster to a P-cluster located at a distance of ∼15.0 Å, bridging the proximal MarDK dimer (Fig. 3b). The P-cluster is symmetrically coordinated by Cys32^MarK^, Cys56^MarK^, Cys121^MarK^, Cys66^MarD^, Cys91^MarD^ and Cys158^MarD^, featuring a centrally shared sulfide (Fig. 2d) and is most likely present in the reduced state (P^N^), as detailed in the Supplementary Discussion 2. Most excitingly, we observed strong electron density at the active site of the methylthio-alkane reductase, located ∼14.4 Å away from the P-cluster (Fig. 2e and Fig. 3b). The density neatly accommodates the [Fe_8_S_9_C]-cluster from Nif(EN)_2_, the FeMoco precursor, missing density for any organic moiety. The lack of (*R*)-homocitrate in Mar(DK)_2_ was confirmed via high-performance liquid chromatography-mass spectrometry (HPLC-MS) (Fig. 1c). To determine the metal composition of these metalloclusters, we performed inductively coupled plasma optical emission spectroscopy (ICP-OES) (Fig. 1b). ICP-OES analysis revealed 32.6 ± 5.3 Fe atoms per catalytic component, consistent with the presence of two P-clusters ([Fe_8_S_7_]-clusters) and two [Fe_8_S_9_C]-clusters per Mar(DK)_2_ heterotetramer, neither Mo nor V were detected. The reductase component MarH_2_ contained 3.5 ± 0.9 Fe atoms, indicating the presence of a labile [Fe_4_S_4_]-cluster. In conclusion, methylthio-alkane reductase is the first Nfl-enzyme described to harbor the large and complex nitrogenase P- and [Fe_8_S_9_C]-clusters.

**Fig. 3:**
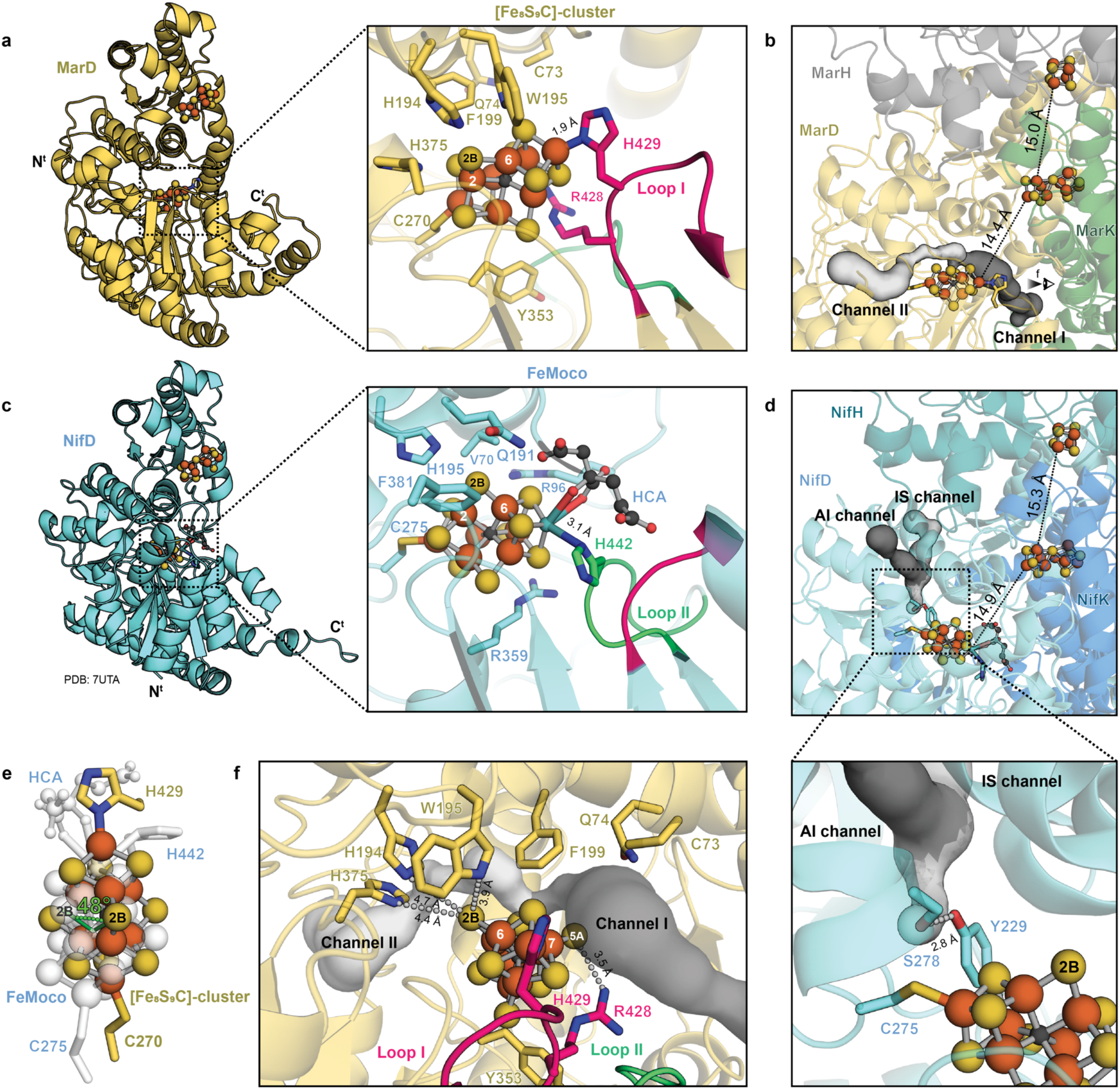
Comparison of active sites and substrate channels from methylthio-alkane reductase and Mo-nitrogenase. **a,c** The active site of the methylthio-alkane reductase (a) features an extended loop (Loop I, highlighted in pink) containing His429^MarD^, which coordinates the [Fe_8_S_9_C]-cluster. Conversely, for Mo-nitrogenase (c; PDB: 7UTA) a different structural domain (Loop II, highlighted in green) contains His442*^Av^*^NifD^, which coordinates FeMoco. Both panels display key amino acid residues potentially involved in metallocluster stabilization and catalysis in stick representation. Metalloclusters and (*R*)-homocitrate (HCA) are shown as ball-and-sticks. N^t^: N-terminus, C^t^: C-terminus. **b,d**, Substrate channels predicted by CAVER^35^ accessing the active sites of the methylthio-alkane reductase (b) and the Mo-nitrogenase (d) are depicted, with the first and second most likely channels colored in dark and light gray, respectively. Mo-nitrogenase channels correspond to the proposed substrate channels by Igarashi and Seefeldt (IS)^32^ and Morrison et. al (AI)^33^. Residues which gate the proximal pocket of the Mo-nitrogenase active site are shown in stick representation^34^. Distances between the three metalloclusters, [Fe_4_S_4_]-cluster, P-cluster and active site cofactor, are indicated by dotted lines. **e**, Overlay of [Fe_8_S_9_C]-cluster and FeMoco from the alignment of MarD and NifD, highlighting the displacement of the S2B belt sulfur position between both clusters. **f**, Side view of the active site of the methylthio-alkane reductase emphasizing S2B and S5A as potential substrate recognition and reduction sites. Distances to S2B and S5A from relevant amino acids are shown as a gray dashed line. Coloring as in Fig. 2a.

### Distinct metallocluster coordination and surrounding

The core of the [Fe_8_S_9_C]-cluster consists of a trigonal prism formed by Fe2-Fe7, which are bridged by three belt sulfides. Assuming a similar role of MarB to NifB in the assembly of the nitrogenase precursor cluster^22^, an interstitial carbide was placed in the center of the cluster, although the resolution of the electron density map is insufficient to locate the carbide. The metallocluster is anchored to the protein scaffold by two amino acid residues, Cys270^MarD^ to Fe1 and His429^MarD^ to Fe8 (Fig. 3a), analogous to the coordination of Fe1 and Mo of FeMoco by Cys275^AvNifD^ and His442^AvNifD^, respectively^23^ (Fig. 3c). However, His429^MarD^ is located at an extended loop specific to MarD, which is not conserved in *bona fide* nitrogenases (Loop I in Fig. 3a,f and Extended Data Fig. 5). This explains why previous sequence analysis of the methylthio-alkane reductase gene cluster had failed to identify this ligand^4^. The octahedral coordination of Mo in the Mo-nitrogenase, which involves three sulfides, (*R*)-homocitrate and His442*^Av^*^NifD^, is replaced in the [Fe_8_S_9_C]-cluster of the methylthio-alkane reductase by a tetrahedral coordination due to the absence of (*R*)-homocitrate. As a result, the anchoring to the protein adopts an axial geometry, and the bond distance between His429^MarD^ and Fe8 of the [Fe_8_S_9_C]-cluster is remarkably shorter (1.9 Å) compared to the bond distance between His442*^Av^*^NifD^ and the Mo atom of FeMoco (3.1 Å in PDB: 7UTA^23^). The observed coordination of the methylthio-alkane reductase [Fe_8_S_9_C]-cluster by just two residues (Cys270^MarD^ and His429^MarD^) indicates that this cluster indeed necessitates a carbide in its center, since it would not be stable without additional ligands.

Beyond the distinct cluster composition, the direct protein environment of methylthio-alkane reductases also deviates from nitrogenases. Substrate binding to nitrogenase metalloclusters has been proposed to occur between Fe2 and Fe6, where the bridging sulfide S2B is replaced by the substrate^24–28^. The conserved residues Val70*^Av^*^NifD^, Gln191*^Av^*^NifD^ and His195*^Av^*^NifD^ surround S2B and are essential for N_2_ reduction^29^ (Fig. 3c). The cavity left by the small side chain of Val70*^Av^*^NifD^allows substrate access to Fe6 of FeMoco^30^. His195*^Av^*^NifD^ forms a hydrogen bond with S2B in the resting state and possibly donates protons during catalysis, while Gln191*^Av^*^NifD^ undergoes structural rearrangements that allow S2B displacement by the substrate during turnover^26,31^. The residues lining the active site of the methylthio-alkane reductase differ significantly (Extended Data Fig. 6a): Cys73^MarD^ and Gln74^MarD^ replace Val70*^Av^*^NifD^, while tryptophan (Trp195^MarD^) and phenylalanine (Phe199^MarD^) residues substitute Gln191*^Av^*^NifD^ and His195*^Av^*^NifD^, respectively (Fig. 3a). However, since the [Fe_8_S_9_C]-cluster of the methylthio-alkane reductase is rotated by ∼48° compared to FeMoco, only Trp195^MarD^ is close enough to S2B to form a hydrogen bond (Fig. 3e,f). Trp195^MarD^, His194^MarD^ and His375^MarD^ surround the S2B site of the [Fe_8_S_9_C]-cluster at distances of 3.9-4.7 Å, and thus could be involved in substrate recognition and binding as well as proton transfer during turnover (Fig. 3a,f). In summary, the methylthio-alkane reductase active site suggests a different substrate recognition and proton transfer to nitrogenases.

### Substrate channels determine specificity

Differences between methylthio-alkane reductase and Mo-nitrogenase extend to substrate channels: Mo-nitrogenase substrate channels proposed by Igarashi and Seefeldt (IS)^32^ and Morrison et. al (AI)^33^ are restricted from accessing the proximal pocket of FeMoco due to hydrogen bond gating by Tyr229*^Av^*^NifD^ and Ser278*^Av^*^NifD^ ^34^ (Fig. 3d). In contrast, the two likeliest methylthio-alkane reductase substrate channels predicted by CAVER^35^ do not show steric impediments. Restricted access to the active site of Mo-nitrogenase could explain its inability to utilize VOSCs, despite being more efficient at reducing smaller substrates (Fig. 1e). The likeliest substrate channel in the methylthio-alkane reductase (Channel I) originates at the interface of MarD and MarK, passes the belt sulfide S5A on the way to S2B and occupies the void left by the absent (*R*)-homocitrate (Fig. 3b,f). The second channel (Channel II) originates at the surface of MarD opposite to channel I and ends at the proposed binding site S2B of the [Fe_8_S_9_C]-cluster (Fig. 3b,f). Besides S2B, belt sulfurs S5A and S3A in FeMoco have been suggested to be involved in N_2_-reduction, since nitrogen-species can also displace these sulfides during turnover^28,36^. In FeMoco, Arg96*^Av^*^NifD^ stabilizes the nitrogen species in position S5A^28^. This arginine is not conserved in methylthio-alkane reductases, instead, its absence contributes to the empty cavity surrounding S5A in Channel I. However, Arg428^MarD^ could potentially fulfill the function of Arg96*^Av^*^NifD^, since it forms a hydrogen bond with S5A (Fig. 3f). Interestingly, Arg428^MarD^ is located at the extended Loop I of MarD, adjacent to the [Fe_8_S_9_C]-cluster coordinating His429^MarD^, and replacing Gly424*^Av^*^NifD^, a residue strictly conserved in all nitrogenases^37^ (Extended Data Fig. 5). Consequently, Arg428^MarD^ together with His429^MarD^ might be a hallmark of methylthio-alkane reductases and distinguish them from nitrogenases. Essentially, the accessibility of S5A and the amino acid environment render it a potential substrate entry site, which could replace S2B in methylthio-alkane reductases.

### The [Fe_8_S_9_C]-cluster does not reduce N_2_

Based on our finding that the methylthio-alkane reductase contains an [Fe_8_S_9_C]-cluster in conjunction with the recent observations that [Fe_8_S_9_C]-cluster-containing nitrogenase maturase Nif(EN)_2_ can reduce N_2_ to NH_3_ to a slight extent^38^, we aimed to determine the N_2_ reduction activity of methylthio-alkane reductases. Surprisingly, no formation of ^15^NH_4_^+^ beyond the background could be detected by nuclear magnetic resonance (NMR) in ^15^N_2_-assays mimicking physiological conditions containing catalytic component Mar(DK)_2_, reductase component MarH_2_, ATP and DT, in contrast to Mo-nitrogenase assays under the same conditions (Fig. 1f and Extended Data Fig. 7). Assays with just the catalytic component Mar(DK)_2_ and the strong reductant europium-II-diethylenetriamine pentaacetic acid (Eu(II)-DTPA) also did not display ^15^NH_4_^+^ formation above background levels (Fig. 1f and Extended Data Fig. 7). This result contrasts with the observation that Nif(EN)_2_ can reduce 1-3 molecules N_2_ in a reductase component- and Eu(II)-DTPA-dependent manner^38^. Thus, our results reveal that the presence of an [Fe_8_S_9_C]-cluster is not the sole determinant for N_2_-reduction, rather the coordination and the immediate surroundings of the [Fe_8_S_9_C]-cluster may determine its activity.

### Electronic structure of metalloclusters

Nitrogenases and Nfl-enzymes exhibit very distinct EPR spectroscopic features^13^, which define the redox state and electronic structure of the metalloclusters. At 4-20 K DT-reduced MarH_2_ exhibited the typical physical spin mixture of a *S=*1/2 (*g=*2.03, 1.933, and 1.84) and a *S=*3/2 species of the [Fe_4_S_4_]^1+^ cluster, characteristic for NifH_2_^39^, of which the relative content of the two spin states is ATP dependent, as detailed in the Supplementary Discussion 3 (Fig. 2f and Extended Data Fig. 8b,c). Next, we analyzed the catalytic component Mar(DK)_2_: upon DT reduction, and in a dye-mediated redox titration with a midpoint potential (E_m_) of −390 mV vs. normal hydrogen electrode (NHE), a relatively sharp *S=*1/2 signal (*g*_average_ = 1.96) appeared (Fig. 2g and Extended Data Fig. 8a). Although the Mar(DK)_2_ properties did not match those of the Mo-nitrogenase Nif(DK)_2_ (see Supplementary Discussion 4; Extended Data Fig. 9 and Extended Data Table 3), it was very similar to the signal assigned to a reduced P-cluster in the catalytic component of V-nitrogenase^40^. Upon oxidation with indigodisulfonate (IDS) or by dye-mediated redox titration with E_m_ = −250 mV vs. NHE a peculiar broad *S=*1/2 signal (simulated with *g*_xyz_ = 1.967, 1.926 and 1.83) with *g*_average_ = 1.91 appeared, totally unlike P^1+^ and P^3+^ EPR signals^13,41^. This EPR signal has previously been observed for the [Fe_8_S_9_C]-cluster-containing nitrogenase maturase Nif(EN)_2_^38^ (Fig. 2g). Our findings show that the *g*_average_ = 1.91 signal is the fingerprint for the active site metallocluster of Mar(DK)_2_ and indicate that a carbide is indeed present in the [Fe_8_S_9_C]-cluster of the methylthio-alkane reductase. By mixing reduced (but DT-free) MarH_2_ with oxidized (but IDS-free) Mar(DK)_2_ and MgATP we could demonstrate that the structurally defined electron transfer pathway (Fig. 2a) guides an electron to the species with *g*_average_= 1.91 (Fig. 2h). Concomitantly, the signal assigned to the reduced P-cluster partially appeared, in agreement with its low redox potential.

### Evolution of nitrogenase metalloclusters

Our structural and biophysical data conclusively show that methylthio-alkane reductases contain metalloclusters until now exclusively associated with nitrogenases. This strongly suggests that their last common ancestor probably contained a P-cluster and a large active site cluster, as previously hypothesized^42^. Using ancestral sequence reconstruction, we inferred the catalytic component sequence of the last common ancestor of nitrogenases and methylthio-alkane reductases and predicted its structure using AlphaFold 2 (Extended Data Fig. 10). In our ancestral sequences, we confidently infer the six cysteines that would ligate a P-cluster. For the residues ligating the [Fe_8_S_9_C]-cluster, we confidently infer a cysteine homologous to Cys270^MarD^, but not a histidine equivalent to His429^MarD^ or His442*^Av^*^NifD^, since this ligand is located in different loops in nitrogenases and methylthio-alkane reductases. Our results indicate that this common ancestor had a P-luster and probably an [Fe_8_S_9_C]-cluster ligated differently than extant nitrogenases and methylthio-alkane reductases. This ancestral reductase was probably catalytically active and contained a complex metallocluster buried in its active site, rather than being a maturase with a surface-exposed cluster like Nif(EN)_2_^15^, although its activity is unresolved, so far. The discovery of nitrogenase clusters in the methylthio-alkane reductase provides a new perspective on how the protein scaffold controls the maturation, insertion, and reactivity of these “great clusters” of biology ^43^ and proves that their history predates the origin of biological N_2_-fixation.

### Conclusions

We aimed to identify the factors determining the unique activities of methylthio-alkane reductases. We revealed that these enzymes contain large nitrogenase metalloclusters, including a P-cluster bridging the MarDK subunits and an [Fe_8_S_9_C]-cluster in the active site of MarD, instead of a [Fe_4_S_4_]-cluster and a substrate-binding site. Consequently, methylthio-alkane reductase is the first nitrogen fixation-like enzyme with such metalloclusters that cannot perform N_2_-fixation. The absence in methylthio-alkane reductase of conserved nitrogenase amino acid residues Val70*^Av^*^NifD^, Gln191*^Av^*^NifD^ and His195*^Av^*^NifD^ involved in N_2_-reduction mechanism^29^ could explain this observation. Moreover, complete nitrogenase activity might require (*R*)-homocitrate, suggested to be crucial for proton transfer during Mo-nitrogenase catalysis^44–46^. While Mo-nitrogenase is likely more reactive and efficient in electron transport, it cannot break the weaker C-S bond of VOSCs reduced by methylthio-alkane reductases. Our structural data suggest that the specificity of methylthio-alkane reductases for methylated sulfur compounds is probably due to wider substrate channels in comparison to nitrogenases, allowing unrestricted access of these larger compounds to the active site. The insights gained in this study highlight the catalytic potential of the [Fe_8_S_9_C]-cluster beyond ammonia formation and open the door to engineering the methylthio-alkane reductase for the biotechnological production of small hydrocarbons such as methane and ethylene.

## Methods

### Chemicals

Chemicals were acquired from Thermo Fisher Scientific (Waltham, Massachusetts, USA), Carl Roth (Karlsruhe, Germany), Sigma-Aldrich (St. Louis, Missouri, USA) and Honeywell (Morristown, New Jersey, USA). Gas bottles were purchased from Air Liquide (Paris, France).

### Bacterial strains and growth conditions

Modified *Rhodobacter capsulatus* B10S^47^ strain MM0246^17^ was used for the heterologous expression and purification of the methylthio-alkane reductase from *R. rubrum*. *R. capsulatus* MM0246 does not encode for a methylthio-alkane reductase^4^. In addition, the Mo-nitrogenase and Fe-nitrogenase genes were disrupted with a spectinomycin or a gentamycin resistance cassette, respectively. *R. capsulatus* was cultured anaerobically under photoheterotrophic conditions (60 W krypton lamps) in modified RCV liquid medium^17^ with the addition of 10 mM serine and 20 µg ml^−1^ streptomycin sulfate.

*Escherichia coli* DH5α (Thermo Fisher Scientific, Waltham, Massachusetts, USA) and *E. coli* ST18^48^ were used for expression plasmid cloning and conjugation into *R. capsulatus* by diparental mating^49^. A list of all strains used in this study is found in Extended Data Table 4.

### Molecular cloning

The *marBHDK* gene cluster (Rru_A0793–Rru_A0796) was amplified from *R. rubrum* DNA by PCR with Q5 high-fidelity DNA polymerase (New England Biolabs, Ipswich, Massachusetts, USA) and cloned under the *R. capsulatus anfH* promoter via Golden Gate assembly^50^ using NEBridge Golden Gate Assembly Kit (New England Biolabs, Ipswich, Massachusetts, USA) into a broad-host range plasmid pOGG024^51^ (pMM0165). To purify the methylthio-alkane reductase subunits separately, affinity tags were added to the sequences via Gibson Assembly^52^ using NEBuilder HiFi DNA Assembly Master Mix (New England Biolabs, Ipswich, Massachusetts, USA; pMM0181). A hexahistidine tag was added to the N-terminus of MarH and a Strep-tag II to the C-terminus of MarD, to purify reductase and catalytic components^53^. The sequence of pMM0165 and pMM0181 were confirmed by whole-plasmid sequencing (Plasmidsaurus, Eugene, Oregon, USA). Primers and plasmids used in this study are listed in Extended Data Tables 5 and 6.

### Anaerobic protein purification and characterization

Purification of Strep-tagged Mar(DK)_2_ and His-tagged MarH_2_ from *R. capsulatus* MM0422 was performed as previously described for the Fe-nitrogenase^17^, except that a modified binding buffer (500 mM NaCl, 50 mM Tris pH 8.5, 10% glycerol, 2 mM sodium dithionite (DT), 0.01% Tween20, 50 mM arginine-glutamate, 20 mM imidazole) was used. Strep-tagged Nif(DK)_2_ and His-tagged NifH_2_ components were purified from *R. capsulatus* MM0480 as previously described^18^. The purity of the proteins was determined by SDS-PAGE, the metal content by inductively coupled plasma optical emission spectroscopy (ICP-OES) and the molecular mass by mass photometry as previously described^17^.

### Homocitrate determination

Semi-quantitative determination of homocitrate was performed using HRES LC-MS. The chromatographic separation was performed on an Agilent Infinity II 1290 HPLC system (Agilent Technologies, Santa Clara, California, USA), using a Kinetex EVO C18 column (150 × 2.1 mm, 3 μm particle size, 100 Å pore size; Phenomenex, Torrance, California, USA) connected to a guard column of similar specificity (20 × 2.1 mm, 3 μm particle size; Phenomenex, Torrance, California, USA) at a constant flow rate of 0.2 ml min^-1^ with mobile phase A being 0.1% formic acid in water and phase B being 0.1% formic acid in methanol at 40°C. The injection volume was 5 µl. The profile of the mobile phase consisted of the following steps and linear gradients: 0 – 0.5 min constant at 0% B; 0.5 – 6 min from 0% to 90% B; 6 – 7 min constant at 90% B; 7 – 7.1 min from 90% to 0% B; 7.1 – 12 min constant at 0% B.

An Agilent 6546 Q-TOF mass spectrometer (Agilent Technologies, Santa Clara, California, USA) was used in negative ionization mode with a dual electrospray ionization source: ESI spray voltage 2000 V, nozzle voltage 500 V, sheath gas 300°C at 11 L min^-1^, nebulizer pressure 45 psig and drying gas 170°C, at 5 L min^-1^. The instrument was used in low molecule size mode with a screening window of 50 - 1700 m/z. 112.9856 and 1033.9881 were used for online mass calibration.

Compounds were identified based on their accurate mass within a mass window of 5 ppm and their retention time compared to standards. Chromatograms were integrated using MassHunter software (Agilent, Santa Clara, CA, USA). Relative abundance was determined based on the peak area by applying a mass extraction window of 10 ppm.

### Methylthio-alkane reductase *in vitro* activity assays

Enzymatic activity of methylthio-alkane reductase and Mo-nitrogenase was determined by *in vitro* activity assays with the substrates (2-methylthio)ethanol (MT-EtOH), ethyl methyl sulfide (EMS), dimethyl sulfide (DMS), protons (under Ar) and acetylene (C_2_H_2_, 10% in Ar) measuring production of ethylene (C_2_H_4_), ethane (C_2_H_6_), methane (CH_4_), molecular hydrogen (H_2_) and C_2_H_4_ as well as C_2_H_6_, respectively. The reductase component (0.15 mg) was added to the catalytic component (0.1 mg) in 700 µl of reaction mix containing 50 mM Tris pH 7.8, 3.5 mM ATP, 7.87 mM MgCl_2_, 44.6 mM creatine phosphate, 0.2 mg ml^−1^ creatine phosphokinase, 5 mM DT and 15 mM of the corresponding VOSC in 1.2 atm Ar, if indicated. Due to the tendency of MarH_2_ to precipitate at high concentrations, all assays were conducted at a catalytic:reductase component molar ratio of 1:5. The reactions took place in 10 ml sealed GC vials, at 30°C and shaking at 200 rpm for 10 min after the addition of the catalytic component. Samples were quenched with 300 µl of 400 mM Na_2_EDTA pH 8.0 up to a total volume of 1 ml. Content of the headspace was determined using a Clarus 690 GC system (Perkin Elmer, Waltham, Massachusetts, USA), as previously described^17^.

### Nuclear magnetic resonance ^15^NH_4_ quantification

*In vitro* nitrogen fixation activity was determined for methylthio-alkane reductase and Mo-nitrogenase by running physiological assays as described above or europium-II-diethylenetriamine pentaacetic acid (Eu(II)-DTPA) assays containing in 1.5 ml volume 5 mg catalytic component Mar(DK)_2_, 20 mM Eu(II)-DTPA, 25 mM Tris pH 8.0 and 2 mM DT. Both assays conditions were performed under 50% ^15^N_2_ (Merck, Darmstadt, Germany) and 50% Ar at 2 atm in 10 ml GC vials, at 30°C, shaking at 250 rpm for 5 h (physiological assays) or 15 h (Eu(II)-DTPA assays) quantifying ^15^NH_4_ with NMR. Negative controls were prepared under Ar or ^14^N_2_ atmospheres. Proteins were removed from the samples by ultrafiltration at 3000 g and 30 min using Amicon Ultra filters (3 kDa cut-off, Merck, Darmstadt, Germany). Subsequently, 500 µl of filtrate were combined with 50 µl of 10 M H_2_SO_4_. 50 µl of d6-DMSO was added as a locking agent to produce a final volume of 600 µl. Two reference samples of 30 mM NH_4_^+^ were prepared using natural abundance ^15^N-NH_4_Cl in assay buffers.

To quantify the concentration of ^14^NH_4_^+^ in each sample, 1D ^1^H spectra with excitation sculpting water suppression were acquired using the zgesgp sequence from the Bruker standard sequence library on an Oxford Instruments 600 MHz magnet equipped with an Avance III console and a TXO Helium cooled cryoprobe. To quantify the concentration of ^15^NH_4_^+^, ^15^N heteronuclear single quantum coherence (HSQC) 1D spectra were acquired on the same hardware using the hsqcetf3gp pulse sequence. 90-degree ^1^H pulse times were calibrated on water. The ^1^H carrier was set on-resonance at water to ensure optimal water-suppression. The ^15^N carrier was set on-resonance for the ammonium signal at 21 ppm. The 90-degree ^15^N pulse time was calibrated on the high signal-to-noise Mo-nitrogenase sample, and assumed constant for the other experiments where calibration was impossible due to low signal-to-noise ratio.

Spectra with varying numbers of scans were recorded for all samples dependent on the observable signal. The signal was rescaled linearly by the number of scans and the receiver gain. The spectra were then processed with zero-filling, an exponential window function and Fourier transformation using the nmrGlue python package^54^.

The ^15^NH_4_^+^ peak was then fitted in the ^15^N HSQC spectrum using a single Lorentzian function within the predefined grey window in Extended Data Fig. 7. The fitted peak was integrated and scaled against the reference peak to provide a ^15^NH_4_^+^ concentration. A region containing only noise was defined between 0 and 1 ppm, and the standard deviation of this region was multiplied by the fitted full-width-at-half-maximum parameter to estimate the peak integral error.

The ^14^NH_4_^+^ peaks were fitted in the water-suppressed spectrum by first finding all local maxima above a spectrum-specific threshold, adding extra manual locations where needed, and fitting the resulting n Lorentzian functions to the data. The ^14^NH_4_^+^ triplet was identified within these fitted peak locations by identifying the characteristic ^14^N-^1^H coupling constant of 52.5Hz. The integrals of the triplet were summed and the fitting error estimated as the standard deviation of the individual triplet peak integrals. These values were scaled against the respective buffer reference samples to provide a concentration of ^14^NH_4_^+^ within the sample.

Two sources of NH_4_^+^ signal were assumed: protein degradation and nitrogen fixation. All protein was produced in unenriched media, so it was assumed to contain a natural abundance 996:4 ratio of ^14^N and ^15^N. Therefore, we assumed that protein degradation contributes a natural abundance ratio of ^14^NH_4_^+^ and ^15^NH_4_^+^ molecules. To account for this, we subtracted a 4/996 fraction of each ^14^NH_4_^+^ concentration from the corresponding ^15^NH_4_^+^ concentration. The subtracted ^15^NH_4_^+^ signal was then used to calculate the total turnover number.

### Cryo-EM sample preparation, collection and processing

MgADP-AlF_3_-trapped Mar(DK)_2_(H_2_)_2_-complex was prepared by starting a 5 ml reaction with 22 nmol of MarH_2_ and 3 nmol of Mar(DK)_2_ in the presence of 100 mM MOPS pH 7.3, 50 mM Tris pH 7.8, 100 mM NaCl, 5 mM DT, 4 mM NaF, 0.2 mM AlCl_3_, 8 mM MgCl_2_ and 1 mM ATP. After 3 h incubation at 30°C and 200 rpm, particles with the appropriate molecular mass were separated via size exclusion chromatography in a HiLoad 26/600 Superdex 200 pg column (Cytiva, Marlborough, Massachusetts, USA) and buffer exchanged to 50 mM Tris pH 7.8, 200 mM NaCl and 5 mM DT. Protein complex with a measured molecular mass of ∼368 kDa (theoretical mass: 344 kDa) was concentrated in Amicon Ultra filters (30 kDa cut-off, Merck, Darmstadt, Germany) to 0.3 mg ml-1. Grids were prepared with a Vitrobot Mark IV (Thermo Fisher Scientific, Waltham, Massachusetts, USA) placed inside an anaerobic tent with a >95% N_2_ <5% H_2_ atmosphere. 4 µl of sample were added to freshly glow-discharged 300 mesh 2/1 µm C-flat copper grids (Electron Microscopy Sciences, Hatfield, Pennsylvania, USA), blotted for 10 s at blot force 4 at 100% relative humidity and 4°C before being plunge frozen in a liquid mix of 37% (v/v) ethane and 63% (v/v) propane.

Cryo-EM data collection was carried out on a Titan Krios G3i electron microscope (Thermo Fisher Scientific, Waltham, Massachusetts, USA) operated at 300 keV in EFTEM mode and equipped with a BioQuantum-K3 imaging filter (Gatan, Pleasanton, California, USA). Data was collected in electron counting mode with EPU (version 3.6) software (Thermo Fisher Scientific, Waltham, Massachusetts, USA) at a nominal magnification of 105,000x, corresponding to a calibrated pixel size of 0.837 Å/pixel, with a total dose of 50 e^−^/Å^-2^ per image, 50 fractions and 2.6 s exposure time. The energy filter slit width was set to 30 eV and the defocus range was −1.0 µm to −2.5 µm at increments of 0.3 µm.

Cryo-EM data analysis was carried out in CryoSPARC v4.4.1^55^ (Extended Data Fig. 3). A total of 16,951 micrographs were subjected to patch motion correction and patch contrast transfer function (CTF) estimation before a template picking using the projections of the Fe-nitrogenase cryo-EM volume as templates (EMD-16890)^17^. An initial set of 10,625,194 particles were extracted at a box size of 380 pixels and used for three consecutive rounds of 2D classification into 200 classes. Particles from the best 2D classes displaying features of at least one MarH_2_ bound to Mar(DK)_2_ were re-extracted at a box size of 380 pixels yielding a total of 3,209,215 particles. These were subjected to ab initio reconstruction into three classes. The best ab initio class with 1,286,797 particles was used for non-uniform refinement yielding a global reconstruction of 2.45 Å with very poor features around both MarD subunits. Hard 3D classification with a local mask on the proximal MarD subunit and 5 Å target resolution resulted in 25 different classes, including classes completely or partially lacking MarD. Particles of the largest class (116,370 particles) displayed the best alignment with reduced particle orientation bias and were subjected to local refinement with a local mask on the proximal MarD and non-uniform refinement, yielding electron density maps with resolutions of 2.86 Å and 2.77 Å, respectively. The final electron density map was obtained by performing a local refinement using a solvent mask yielding a final resolution of 2.75 Å. The resulting map was autosharpened in Phenix v1.21.1^56^. Details on cryo-EM data collection, model building, and refinement statistics can be found in Extended Data Table 1.

### Model building and refinement

The cryo-EM map was initially fitted in ChimeraX-1.7.1^57^ into an AlphaFold 2^58^ multimer model of the Mar(DK)_2_H_2_-complex. Manual refinement and cofactor building was performed in Coot v0.8.9.2^59^. Automatic model refinement was performed in Phenix v1.21.1^56^.

### Substrate channel estimation

Substrate access channels to the active site of the methylthio-alkane reductase were calculated using the tool CAVER Analyst 2.0 BETA^35^. The starting point for channel calculating was the proposed binding site S2B of the [Fe_8_S_9_C]-cluster. The probe radius, shell radius and shell depth were set to 0.6, 3.0, and 4.0 Å, respectively. The two most likely substrate channels, with a maximum bottleneck radius of 27.489 Å and 25.645 Å, respectively, were selected for analysis (Fig. 3b,f).

### EPR spectroscopy

All EPR samples were prepared under anaerobic conditions in a Coy anaerobic glove box. All protein samples were buffer exchanged with Sephadex G-25 packed PD-10 desalting columns (Cytiva, Marlborough, Massachusetts, USA) into 500 mM NaCl, 50 mM Tris pH 8.5 and 10% glycerol to remove excess DT prior to sample preparation.

For the redox titrations, samples were titrated in the presence of a cocktail of 40 µM of each redox mediator through the addition of different volumes of buffered DT or potassium ferricyanide solutions (1.5-100 mM) until the desired redox potential was reached. The mediator mix consisted of 2,6-dichlorophenolindophenol (E_0_=+217 mV), phenazine methosulfate (E_0_=+55 mV), methylene blue (E_0_=+11 mV), resorufin (E_0_=-51 mV), 5,5’-indigodisulfonate (IDS, E_0_=-125 mV), 2-hydroxy-1,4-napthaquinone (E_0_=-152 mV), sodium anthraquinone 2-sulfonate (E_0_=-225 mV), phenosafranin (E_0_=-252 mV), safranin O (E_0_=-289 mV), neutral red (E_0_=-329 mV), benzyl viologen (E_0_=-358 mV) and methyl viologen (E_0_=-449 mV). Upon stabilization of the redox potential, a 300 µl sample was withdrawn, transferred to an EPR quartz tube, capped and shock frozen in liquid nitrogen. The InLab Argenthal microelectrode (Ag/AgCl, E=+207 mV versus H_2_/H^+^, with combined Pt counter electrode) was calibrated with a quinhydrone saturated pH 7 reference buffer solution (E=+285 mV versus H_2_/H^+^ at 25°C).

MarH_2_ samples were prepared by gentle addition of anaerobic stock solutions of the compounds of interest (2 mM DT, 5 mM MgATP, final concentration) to the protein, incubated for at least 3 minutes and frozen as for the redox titration. For the turnover experiment Mar(DK)_2_ was oxidized by addition of 2 equivalents IDS and incubated for 15 min. DT-reduced MarH_2_ and the IDS-oxidized Mar(DK)2 were desalted. Directly after desalting 300 µl samples of MarH2 and oxidized and Mar(DK)_2_ were frozen separately. DT-removed MarH_2_ and IDS-removed Mar(DK)_2_ were mixed in ratio of 1.2:1 reductase:catalytic component with MgATP (5 mM final concentration) incubated for 1 min, concentrated with an Amicon Ultra filter (100 kDa cut-off, Merck, Darmstadt, Germany) to a final volume of 300 µl and shock frozen. Nif(DK)_2_ was purified as described^18^, desalted and treated with 2 mM DT, 0.5 mM IDS or 50 µM IDS.

Cw-EPR spectra were recorded with a commercial Bruker spectrometer composed of an X-band E580-10/12 bridge and a 4122HQE resonator, for regular (perpendicular mode, for Fig. 2 and Extended Data Fig. 8) and an ER4116 dual mode cavity for parallel mode (for Nif(DK)_2_ and trials to detect *g=*12 or *g=*16 signals in Mar(DK)_2_) measurements. The system was equipped with an Oxford Instruments temperature controller and ESR 900 cryostat, which was cryocooled by a Stinger (Cold Edge Technologies, Allentown, Pennsylvania, USA) linked to helium compressor (Sumitomo F-70). Spectral simulations were performed with GeeStrain5^60^ or in Excel (Gaussian curves representing the absorption-shaped highest *g*-value of the very anisostropic doublets of the *S=*3/2 system of MarH_2_, or the first derivative of a Gaussian curve for the isotropic *g=*4.3 EPR signal of the middle doublet of high spin ferric iron). Except for Fig. 2f and Extended Data Fig. 9, five point moving averages were used to limit the noise of EPR spectra.

### Phylogenetic analysis and ancestor prediction

For the phylogeny in Extended Data Fig. 10, sequences from archaeal *cfbD*, *R. rubrum marD*, and *A. vinelandii* nitrogenase operon *nifDKEN* were employed as queries across major archaeal and bacterial lineages. Discrimination between various nitrogenase components was achieved through reciprocal best-blast hits and manual verification of synteny. Subsequently, sequences from both data sets were aligned using Muscle v5^61^ and further refined using clipKIT^62^. The resulting phylogenetic trees were constructed using Iqtree2^63^ under the LG+G4 model, with 35,000 ultrafast bootstraps and 1500 SH-aLRT replicates^64^. Model selection for evolution was based on the Corrected Akaike information criterion (-m MFP -merit AICc option in Iqtree2). The resulting tree was constrained to have *bchlYN* and *bchlBZ* clades sister to *nifDE-likeD* and *nifKN-likeK*, respectively. Every other node was polytomized. The resulting ML analysis of the constrained tree shows similar likelihood values as for the unconstrained tree and it was not rejected by any statistical test performed with iqtree. This topology was chosen for analysis due to the more parsimonious history in terms of subunit oligomerization. Supporting values for this topology were calculated with 500 Feslestein bootstrap analysis using Iqtree2 and the approximate likelihood ratio test (aLRT) using PhyML. Ancestral sequence reconstruction for the nodes of interest was performed using a custom script that uses Iqtree2 and PAUPv4 using the LG+G4 model.

### AlphaFold Structure Prediction

The structure of the last common nitrogenase and methylthio-alkane reductase ancestor AncD and AncK was predicted by AlphaFold 2^58^ using ColabFold^65^ with default parameters (without templates, model_type: alphafold2_multimer_v3, num_relax: 1).

## Data and materials availability

The refined MarDK_2_H_2_ structure and electron density map were deposited under PDB-ID 9FMG and EMD-50553, respectively. Raw data of SDS-PAGE analysis, ICP-OES and mass photometry measurements, gas chromatography and NMR enzyme activity assays, EPR spectroscopy and protein sequences used in multiple sequence alignments are deposited on Edmond, the Open Research Data Repository of the Max Planck Society (https://doi.org/10.17617/3.OHDUAZ), which will be released upon acceptance. All NMR data and the python package created for analysis have been uploaded to GitHub and can be found here: https://github.com/charliebuchanan/nitrogenaseNMR.

## Acknowledgments

The authors thank the Central Electron Microscopy Facility at the Max Planck Institute of Biophysics for expertise and access to their instruments. We thank C. Thölken, P. Klemm and M. Lechner for assistance with data transfer and computing cluster access. We thank V. Reitz for her help with molecular cloning. We thank C. Scholz for access and measurements of ICP-OES. We thank L. Ernst for his help with biochemistry work and purification of the Mo-nitrogenase. We thank F. Hagen for access to the simulation program GeeStrain5. We thank Y. Hu and M. Ribbe for providing an EPR spectrum of [Fe_8_S_9_C]-cluster-bound Nif(EN)_2_.

## Funding

This work was supported by the German Research Foundation (grant 446841743, J.G.R). A.L.M., J.Z., C.J.B., N.P., G.K.A.H and J.G.R. are grateful for generous support from the Max Planck Society. A.L.M. acknowledges the financial support by the International Max Planck Research School Principles of Microbial Life. The BBSRC (BB/R000255/1) is acknowledged for supporting the NMR facility at University College London. The EPR spectrometer upgrade and closed-cycle cryostat (A.J.P.) was funded by the German Research Foundation (248/320-1, project number 444947649) and the Rhineland-Palatinate government. A.J.P. acknowledges Silke Leimkühler and the SPP1927 FeS for Life (grant PI610/2-1 and 2). This research is supported by the UKRI and EPSRC (DFH; EP/X036782/1). G.T.H. was supported by a BBSRC Discovery Fellowship (BB/X009955/1). C.J.B is grateful for the support of a Human Frontiers Science Programme Long Term Fellowship (LT0018/2024-L). Open Access publishing was enabled and organized by Projekt DEAL.

## Author contributions

J.G.R. conceived and supervised the project. J.G.R. acquired funding. A.L.M. and J.G.R. planned and analyzed experiments. A.L.M. performed molecular work, anaerobic protein purification, enzyme biochemistry, mass photometry measurements and analysis of the structural data. F.V.S. performed anaerobic Mo-nitrogenase purification and activity assays. S.L. and G.K.A.H. performed the phylogenetic analysis. N.P. performed homocitrate measurements. C.J.B., G.T.H. and D.F.H. performed NMR measurements and analysis. A.L.M. and S.P. performed cryo-EM data acquisition. A.L.M., J.Z. and T.R.T. processed and refined the cryo-EM structure. A.L.M., J.S. and A.J.P. performed EPR measurements and analysis. A.L.M., J.G.R., J.S. and A.J.P. wrote the original manuscript which was reviewed and edited by all authors.

## Competing interests

The authors declare no competing interests.

## Additional Information

Supplementary Information is available for this paper.

## Extended Data

**Extended Data Fig. 1:**
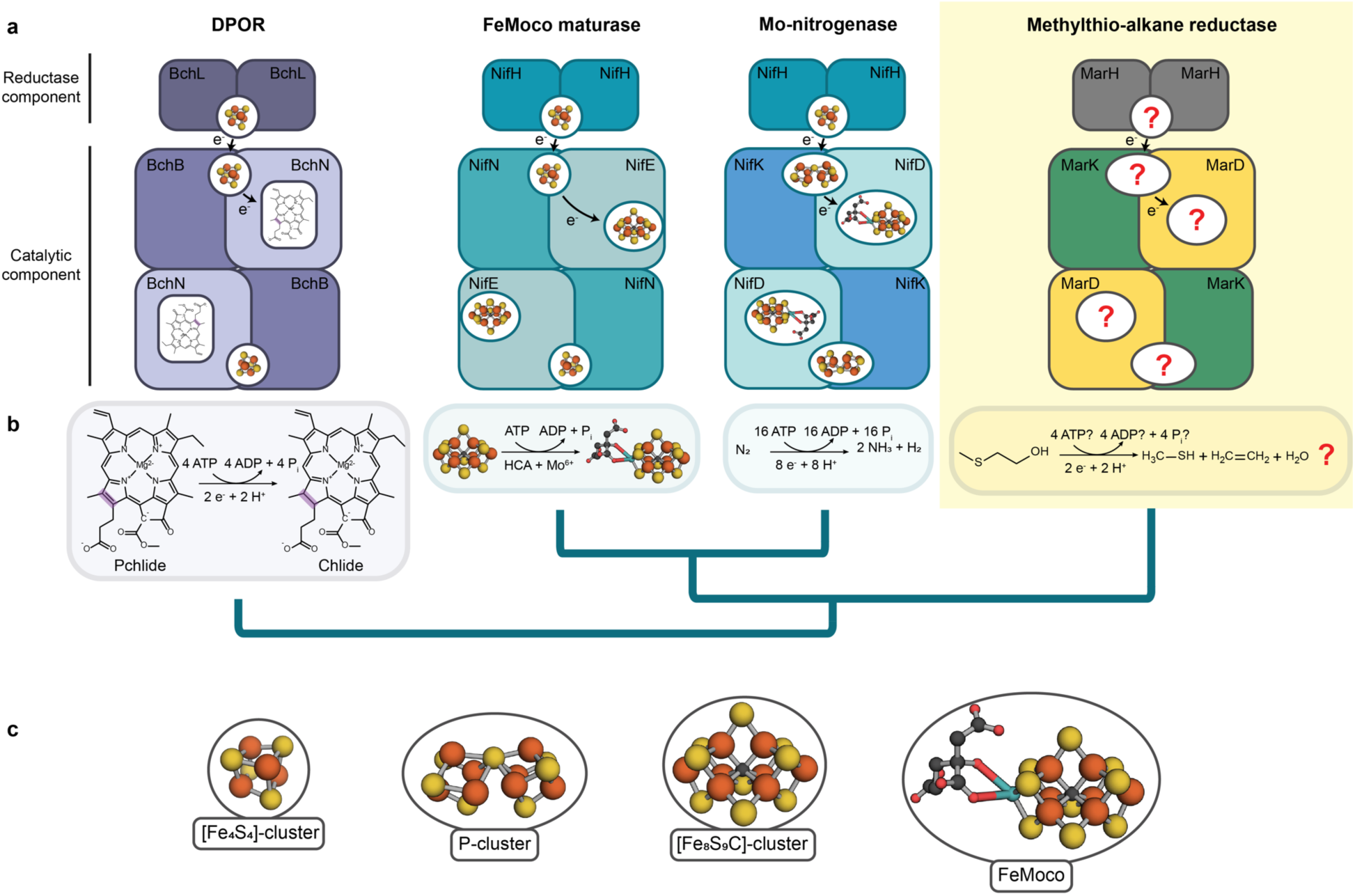
Structural and functional diversity of the nitrogenase and nitrogen fixation-like (Nfl) protein family, highlighting the methylthio-alkane reductase characterized in this study. **a**, Schematic overview illustrating structures from key nitrogenase and Nfl proteins with their associated metalloclusters responsible for electron transfer and catalysis. Electrons are sequentially transferred from the [Fe_4_S_4_]-cluster of the reductase components to the metal cofactors of the catalytic components. For DPOR and FeMoco maturase Nif(EN)_2_, this is a [Fe_4_S_4_]-cluster^15,66^, while Mo-nitrogenase uses a P-cluster ([Fe_8_S_7_]-cluster) as an electron relay to the active site^11^. In DPOR, the substrate protochlorophyllide (Pchlide) sits directly at the active site^66,67^. Nitrogenases harbor more complex metalloclusters in their active sites, such as the FeMoco^11^, Nif(EN)_2_ harbors the [Fe_8_S_9_C]-cluster^15^. The metallocluster composition for the methylthio-alkane reductase as well as the stoichiometry of the reaction was still unresolved. **b**, Main catalytic reactions. DPOR reduces the C17=C18 double bond of Pchlide to chlorophyllide *a* (Chlide) in the chlorophyll *a* biosynthetic pathway^66,67^. The Nif(EN)_2_ maturase converts the precursor [Fe_8_S_9_C]-cluster into the FeMoco by inserting Mo and (*R*)-homocitrate^15^. Mo-nitrogenase reduces protons and N_2_ to NH_3_ and H_2_^11^. The methylthio-alkane reductase is proposed to reduce MT-EtOH to methanethiol and C_2_H_4_ ^4^. Protein phylogenetic relationships are depicted based on fig. S10. **c**, Legend of the metalloclusters found in nitrogenase and Nfl proteins shown in (a).

**Extended Data Fig. 2:**
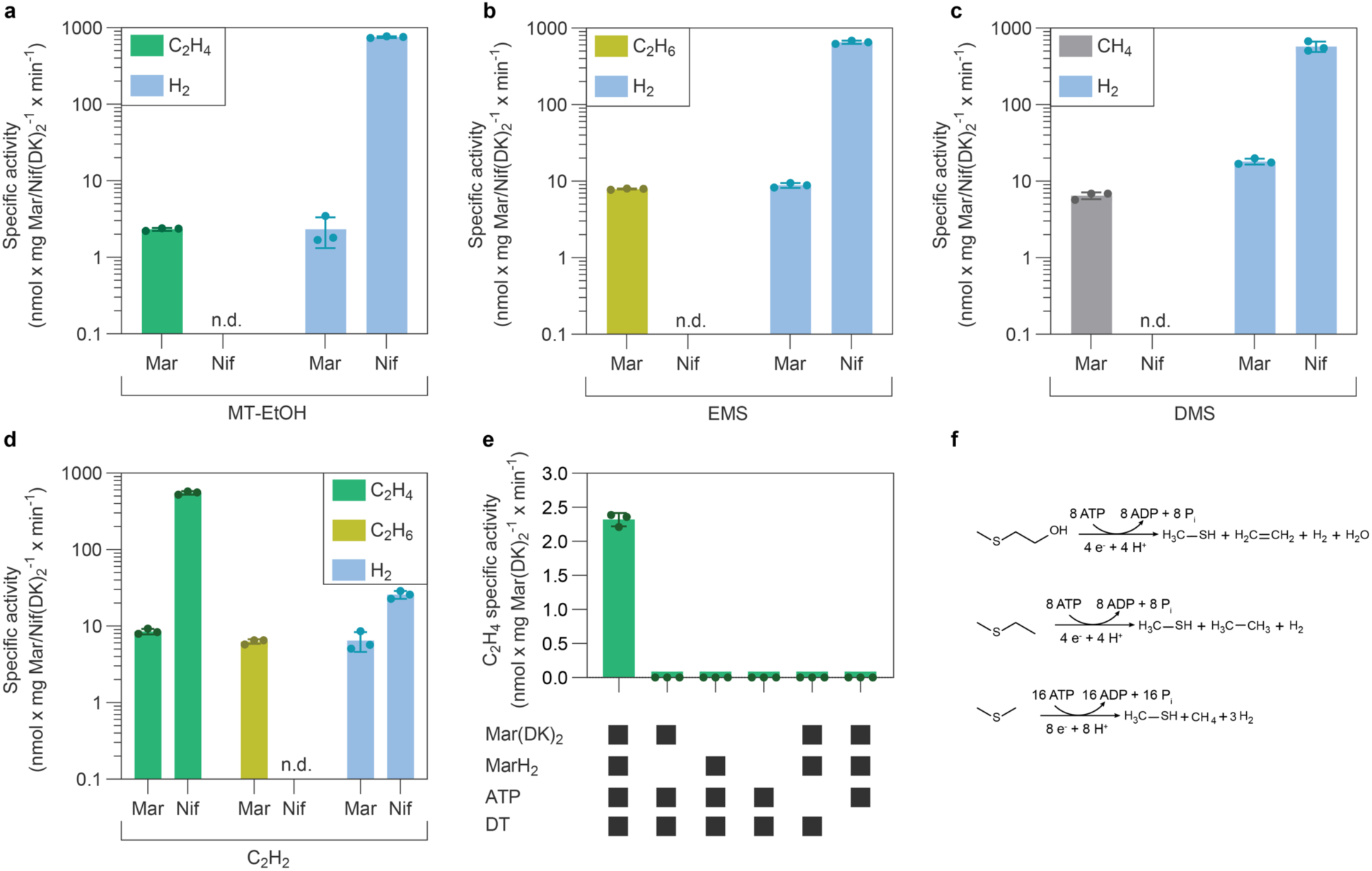
Substrate reduction and product formation of methythio-alkane reductase and Mo-nitrogenase. **a-d**, *In vitro* activities of methylthio-alkane reductase (Mar) and Mo-nitrogenase (Nif) for the reduction of (2-methylthio)ethanol (MT-EtOH, a), ethyl methyl sulfide (EMS, b), dimethyl sulfide (DMS, c) and acetylene (C_2_H_2_, d). Specific activities are plotted for the formation of ethylene (C_2_H_4_, green), ethane (C_2_H_6_, yellow), methane (CH_4_, gray) and hydrogen (H_2_, blue) (n=3). n.d., not detected. **e**, *In vitro* C_2_H_4_ specific activity of the methylthio-alkane reductase for the reduction of MT-EtOH is dependent on the addition of reductase and catalytic components, ATP and DT (n=3). **f**, Stoichiometric reactions for the reduction of MT-EtOH, EMS and DMS by the methylthio-alkane reductase.

**Extended Data Fig. 3:**
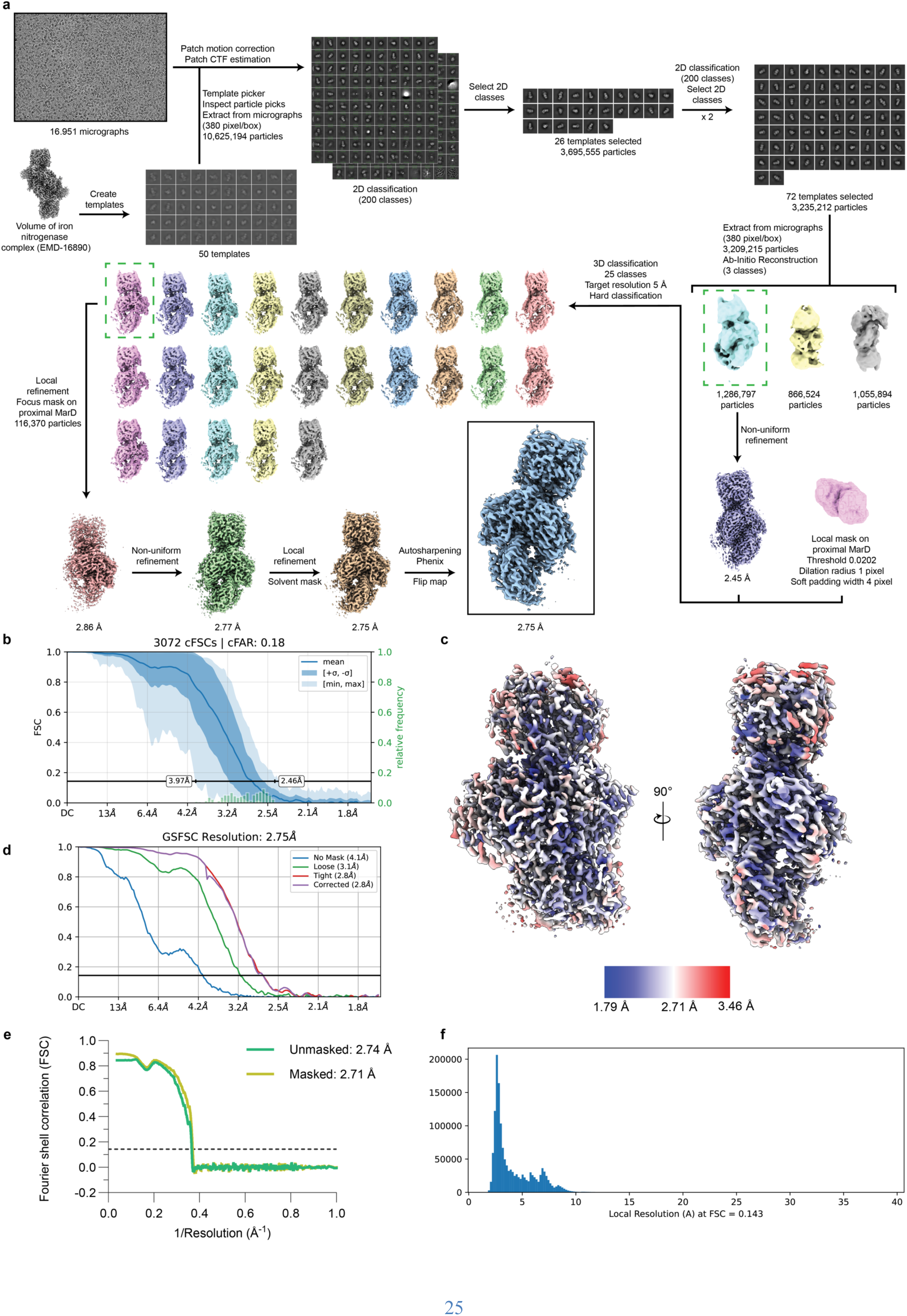
Cryogenic electron microscopy data acquisition and analysis of the methylthio-alkane reductase complex. **a**, Schematic of the processing workflow for cryo-EM data using CryoSPARC v4.4.1^55^ and Phenix v1.21.1^56^. **b**, Conical FSC Area Ratio (cFAR) plot by CryoSPARC v4.4.1 assessing specimen orientation bias. **c**, Local resolution estimated by CryoSPARC v4.4.1 mapped onto the final electron density map contoured at level 10. **d**, Gold-Standard Fourier Shell Correlation (GSFSC) plot by CryoSPARC v4.4.1 with resolution calculated at Fourier shell correlation (FSC)=0.143. **e**, Map to atomic model FSC plot determined at FSC = 0.143. **f**, Histogram of local resolution values calculated at FSC=0.143 by CryoSPARC v4.4.1.

**Extended Data Fig. 4:**
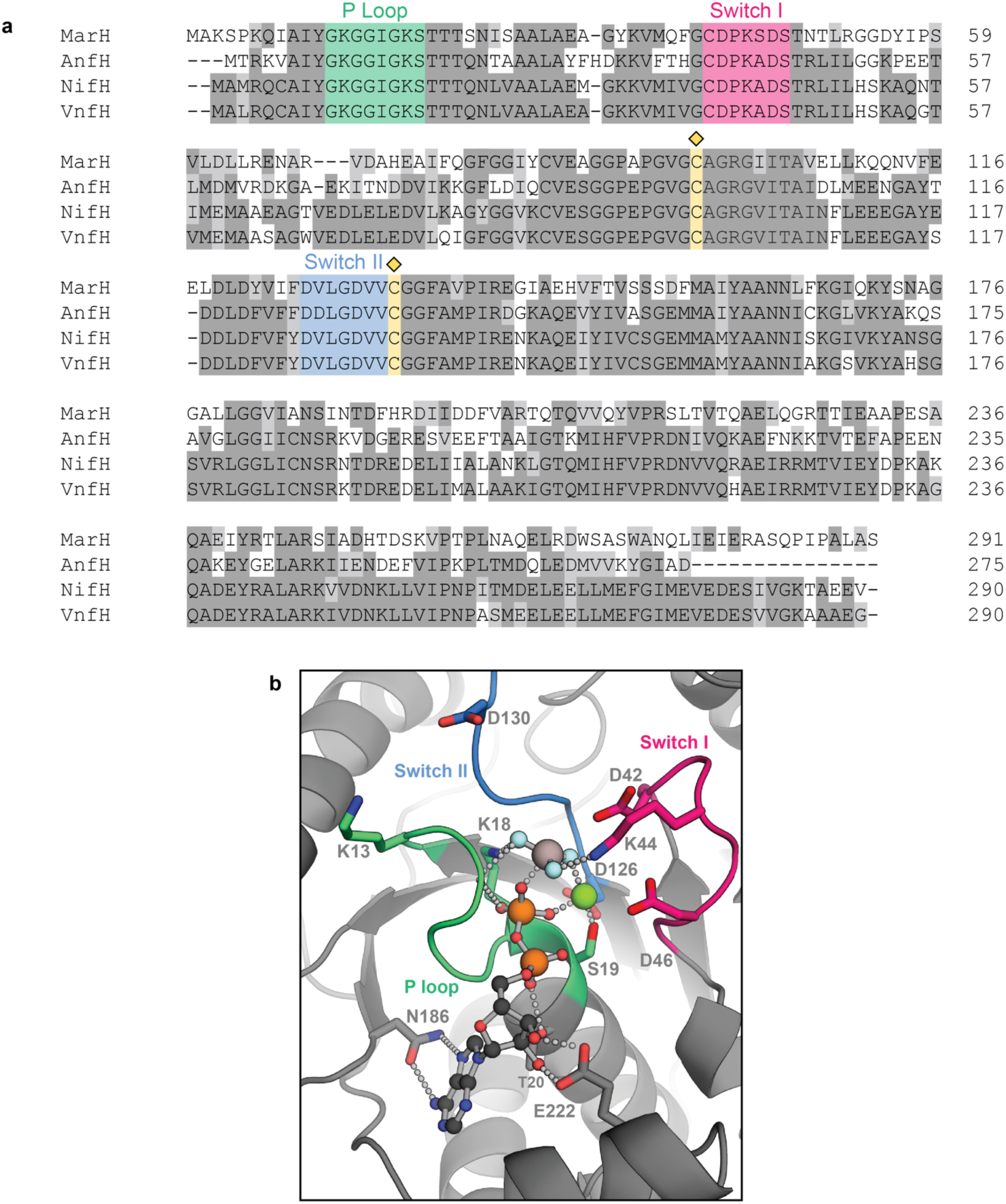
The reductase component MarH_2_ belongs to the family of P-loop NTPases. **a**, Clustal Omega^68,69^ multiple sequence alignment (MSA) of the *R. rubrum* methylthio-alkane reductase reductase component MarH to the *A. vinelandii* Mo-, V- and Fe-nitrogenase reductase components NifH, VnfH and AnfH, colored with Boxshade v3.3. The conserved P-loop, switch I and switch II regions involved in nucleotide binding and hydrolysis are colored in green, pink and blue, respectively. Cys97^MarH^ and Cys133^MarH^ coordinating the [Fe_4_S_4_]-cluster are highlighted in yellow. **b**, Close-up view of the MgADP-AlF_3_ moiety bound to one MarH subunit. P-loop, switch I and switch II regions are colored as in (a). MgADP-AlF_3_ is represented in ball-and-stick style with carbon atoms colored in dark gray, nitrogen in blue, oxygen in red, phosphorus in orange, aluminium in light gray, fluoride in pale cyan and magnesium in green. Amino acid residues coordinating the ligand by hydrogen bonding are shown in stick representation.

**Extended Data Fig. 5:**
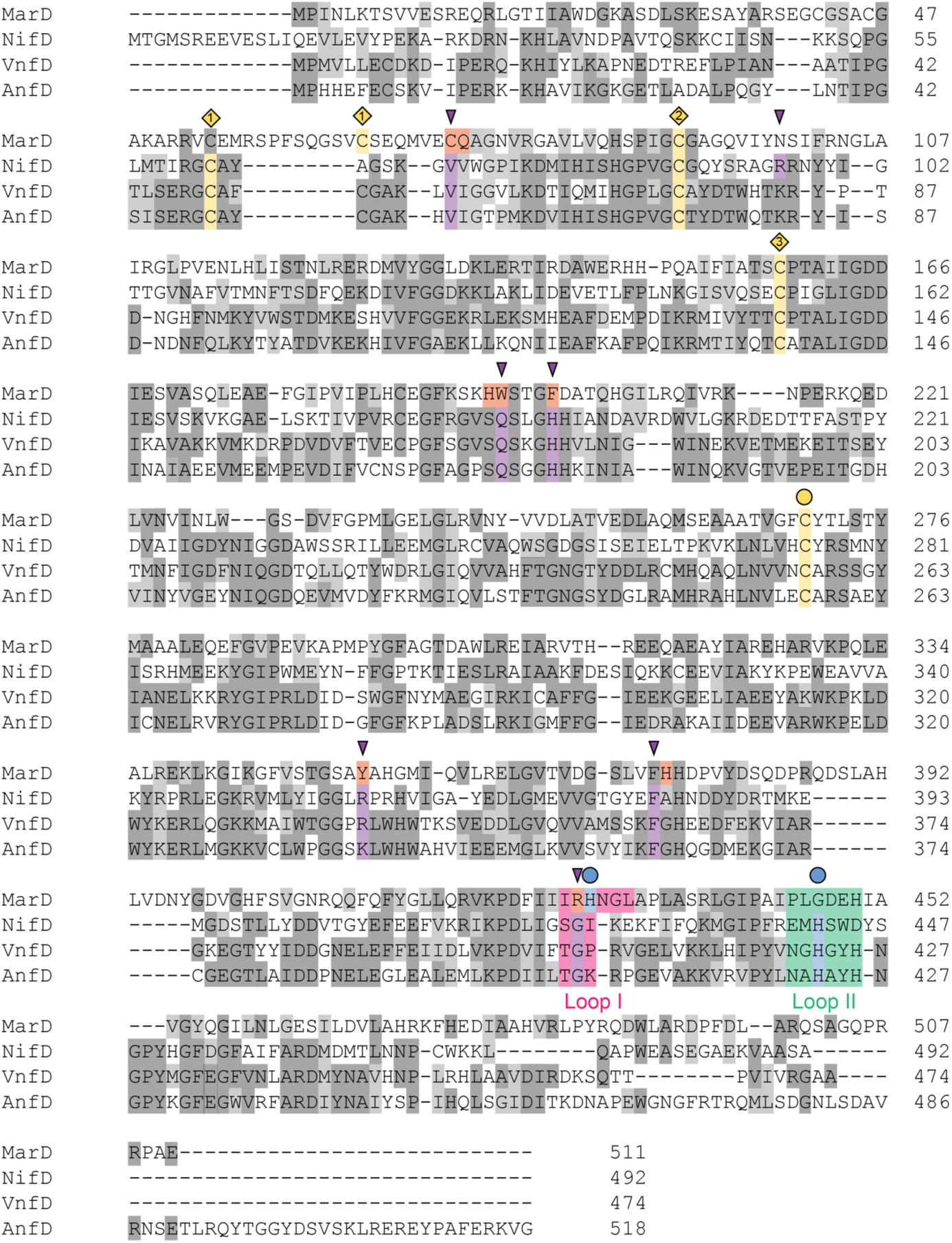
Sequence alignment of the catalytic D-subunits of methylthio-alkane reductase and Mo-, V-, and Fe-nitrogenase. Clustal Omega^68,69^ MSA of the *R. rubrum* methylthio-alkane MarD subunit and the *A. vinelandii* Mo-, V- and Fe-nitrogenase NifD, VnfD and AnfD subunits, colored with Boxshade v3.3. Cysteine triads involved in P-cluster coordination are highlighted in yellow and numbered 1-3. Cysteine and histidine ligands responsible for the coordination of the [Fe_8_S_9_C]-cluster, FeMoco, FeVco and FeFeco are labeled with a yellow and blue circle, respectively. Strictly conserved residues involved in nitrogenase catalytic mechanism and FeMoco, FeVco and FeFeco stability are highlighted in purple, while residues replacing them in the structure of the methylthio-alkane reductase are highlighted in orange. The extended loop I of the methylthio-alkane reductase containing the coordinating histidine His429^MarD^ is highlighted in pink, while the Mo-nitrogenase loop II containing the coordinating histidine His422*^Av^*^NifD^ is highlighted in green.

**Extended Data Fig. 6:**
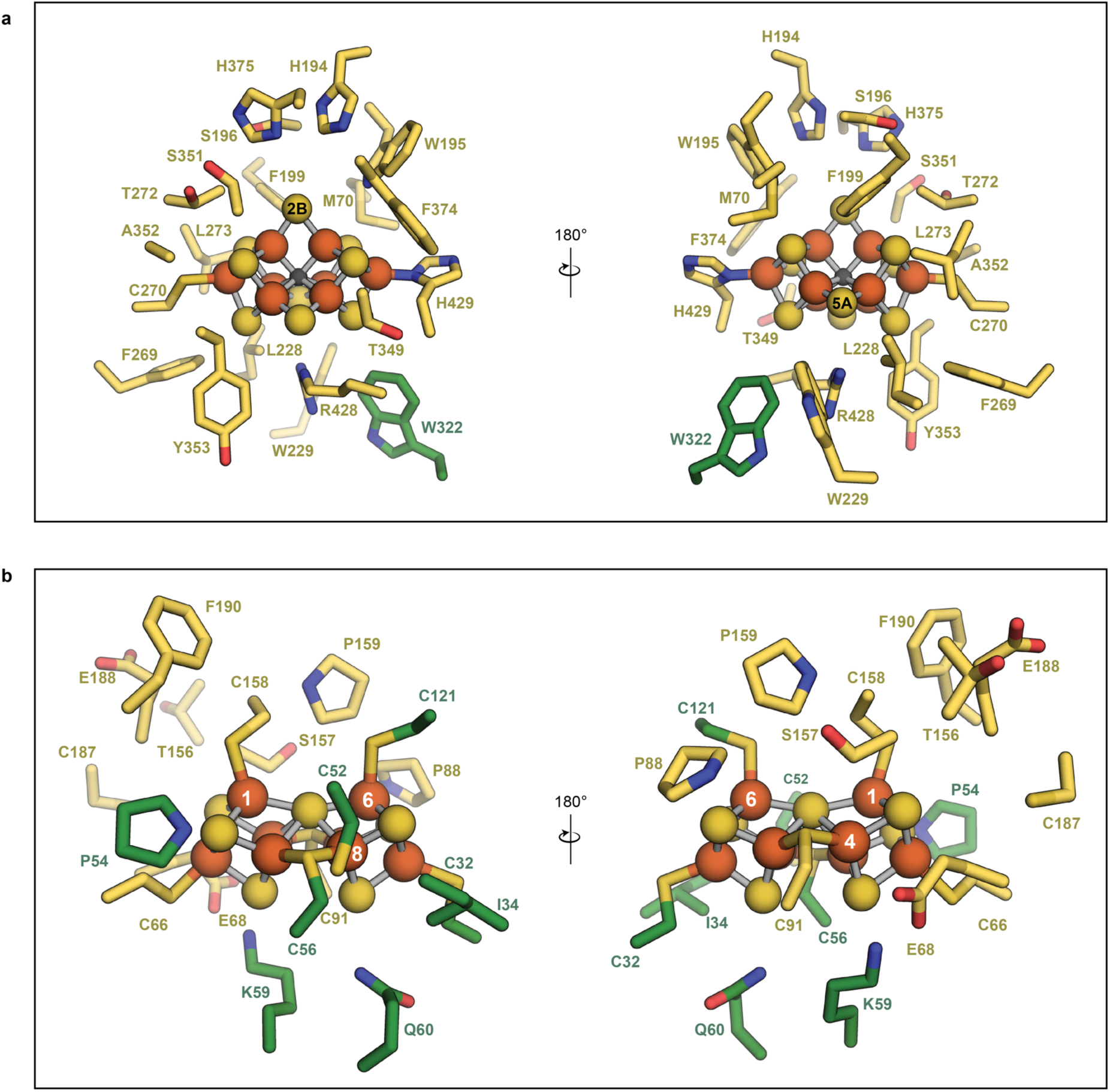
Environment of the methylthio-alkane reductase [Fe_8_S_9_C]-cluster and P-cluster. **a,b**, Close-up view of the [Fe_8_S_9_C]-cluster of the active site of the methylthio-alkane reductase (a) and the P-cluster bridging the MarDK subunits (b). Metalloclusters are shown as ball-and-sticks, with carbon atoms in dark gray, sulfur in yellow, and iron in dark orange. Amino acid residues surrounding the metalloclusters within a 5 Å radius are shown in stick representation. Amino acid residues of MarD and MarK are colored in yellow and green, respectively.

**Extended Data Fig. 7:**
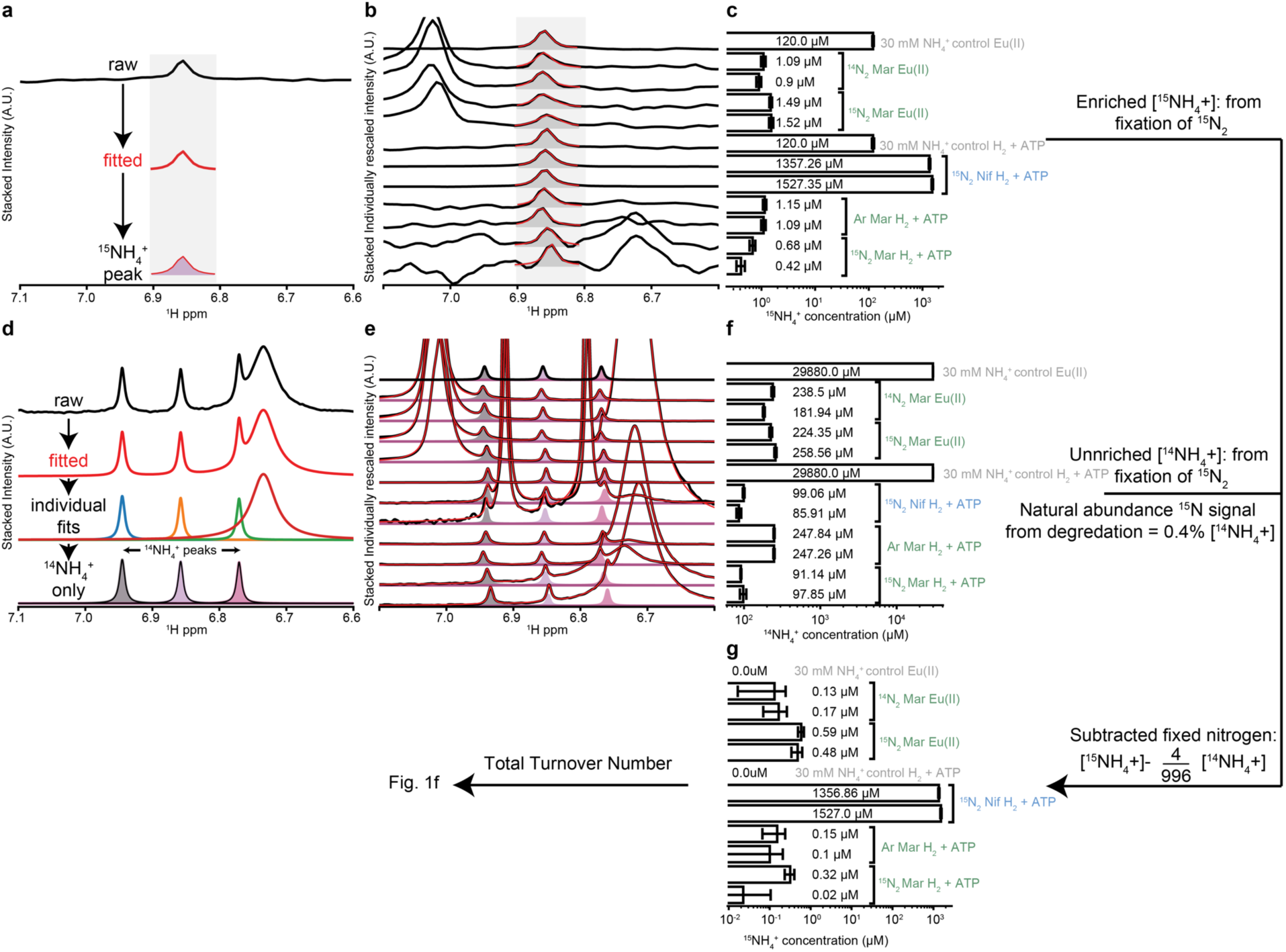
Nitrogen assays with methylthio-alkane reductase and product detection by nuclear magnetic resonance spectroscopy. A single Lorentzian function is fitted (**a**, red) to each processed NMR spectrum from the ^15^N heteronuclear single quantum coherence experiment (**a**, black), before being integrated (**a**, pink shaded) and scaled against reference NH_4_^+^ samples to provide a concentration. Performing this on all samples (**b**) shows good agreement between fitted (**b**, red) and raw (**b**, black) data. The extracted ^15^NH_4_^+^ concentrations span the nanomolar to millimolar range (**c**). n Lorentzian functions are fitted (**d**, red and individual fits) to the processed water-suppressed ^1^H NMR data and the ^14^NH_4_^+^ peaks are extracted by identifying the fitted peak locations split by the characteristic 52.5 Hz coupling constant (**d**, shaded). The summed integral of the triplet is then scaled against reference NH_4_^+^ samples to provide a concentration. This was performed on all samples and the resulting fits (**e**, red) show good agreement with raw (**e**, black) data. The ^14^NH_4_^+^ concentrations span the 100 µM to 250 µM range for the protein samples (**f**). As the protein is isotopically non-enriched, we assume the degradation products will have the natural abundance distribution of 996:4. To account for this, we subtract 0.4% of the ^14^NH_4_^+^ concentration from the ^15^NH_4_^+^ concentration to produce a subtracted ^15^NH_4_^+^ concentration that more faithfully describes the maximum ^15^NH_4_^+^ concentration that could have arisen from nitrogen fixation (**g**). The data was used to calculate the total turnover numbers shown in Fig. 1f.

**Extended Data Fig. 8.**
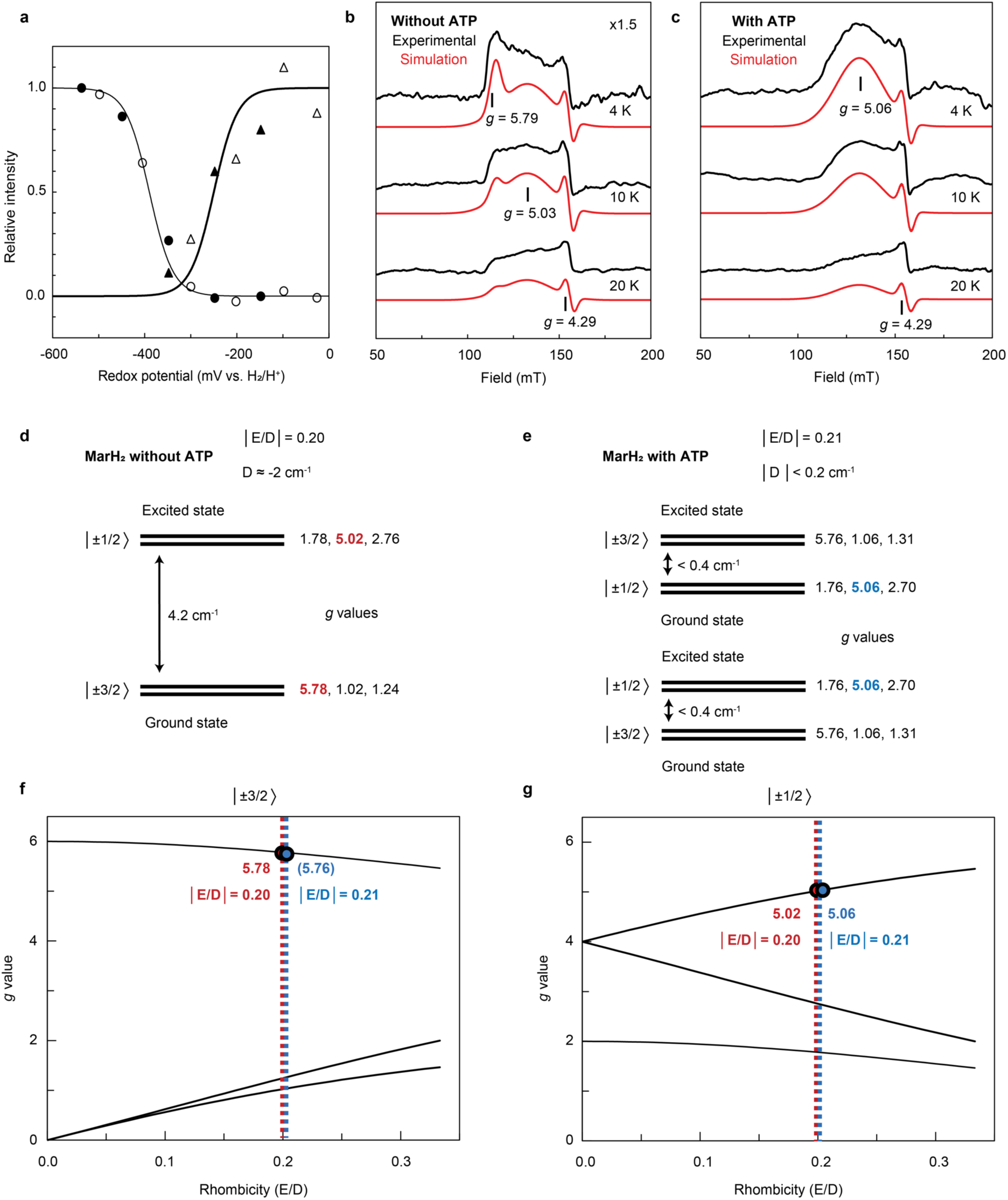
EPR spectroscopy of Mar(DK)_2_ and MarH_2_ metalloclusters. **a**, Amplitudes of the two species from Fig. 2g and further redox titrations of Mar(DK)_2_ with fits to the Nernst equation for n=1 with E_m_ = −390 and −250 mV. **b,c**, *S=*3/2 EPR signals of the [Fe_4_S_4_]^1+^ cluster in dithionite (DT)-reduced MarH_2_ with (c) and without (b) MgATP. EPR signals recorded at 4, 10 and 20 K in black lines, and with simulations in red lines (parameters in Extended Data Table 2). EPR conditions: microwave frequency 9.35 GHz; modulation frequency, 100 kHz; 1.5 mT modulation amplitude; microwave power, 20 mW. **d,e**, Energy level diagrams of the *S=*3/2 spin system of MarH_2_. The calculated *g*-values for MarH_2_ without ATP (d) are in red, those for MarH_2_ in the presence of ATP in blue (e). The intensity of the *g=*5.06 peak shows Curie law behaviour, indicating that both doublets are almost equally populated. Thus, it could not be determined whether the peak is from the ground or excited state, and the magnitude and sign of the D value could not be estimated. The two possible energy level diagrams are shown. **f,g**, Rhombograms for the two |±3/2) and |±1/2) doublets of the *S=*3/2 state^70^. Calculated g-values as function of the rhombicity (E/D) are shown.

**Extended Data Fig. 9.**
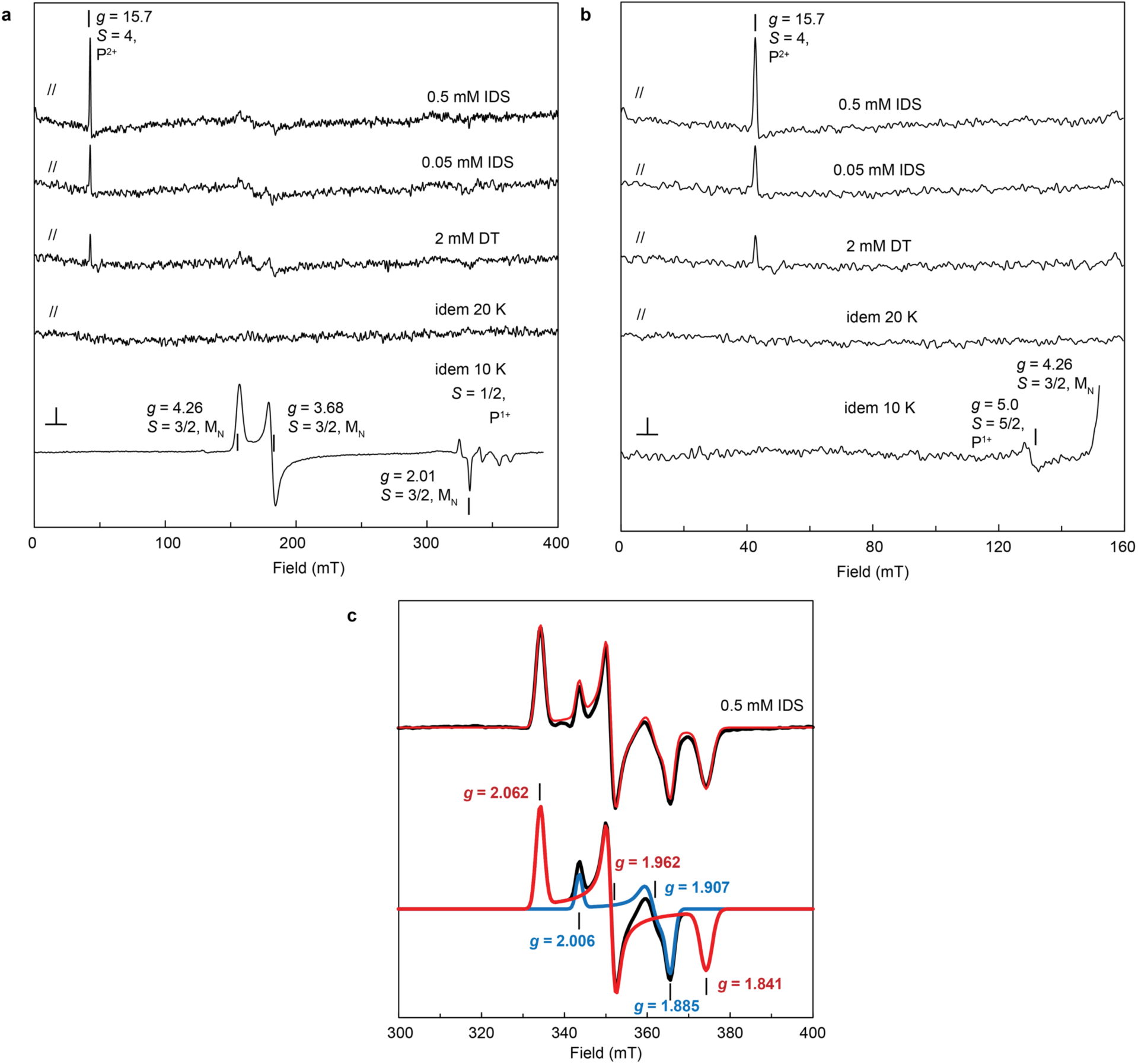
EPR spectroscopy of the metalloclusters of *Rhodobacter capsulatus* Nif(DK)_2_. **a**, Samples of desalted Nif(DK)_2_ were treated with indicated oxidants (5,5’-indigodisulfonate, IDS) or reductant (dithionite, DT) and measured at 4 K (or as indicated otherwise). The strong *S=*3/2 EPR signal in DT-reduced Nif(DK)_2_ (bottom trace, x0.1 in comparison to the other traces) was accompanied by a weak *S=*1/2 EPR signal in the measurement with the perpendicular mode of a dual mode cavity. These signals disappear in the parallel mode. However, at 4 K a very sharp *g=*15.6 signal of the *S=*4 state of the P^2+^ cluster was detected, which increased several fold upon progressive IDS oxidation. The signal is from an excited state, as its intensity decreased with increased temperature (see trace at 20 K). **b**, Zoom in of (a), with the bottom trace on the same scale as the other traces to reveal a weak derivative-shaped *g=*5.0 signal belonging to the middle g-value of the |±1/2) doublet in the *S=*5/2 state of the P^1+^ cluster. The corresponding broad *g=*6.7 peak could not be detected. **c**, The *S=*1/2 EPR signal at 20 K of the P^1+^ cluster in 0.5 mM IDS-oxidized Nif(DK)_2_, which could be simulated by a mixture of two species. Top traces: experimental spectrum (black line) with the sum of the simulated spectra (red line, parameters in Extended Data Table 3). Bottom traces: individual simulations in red and blue lines, together with their sum (black line). EPR conditions: microwave frequency, 9.63 GHz (perpendicular mode) or 9.34 GHz (parallel mode); modulation frequency, 100 kHz; 0.5 mT modulation amplitude, microwave power, 20 mW.

**Extended Data Fig. 10:**
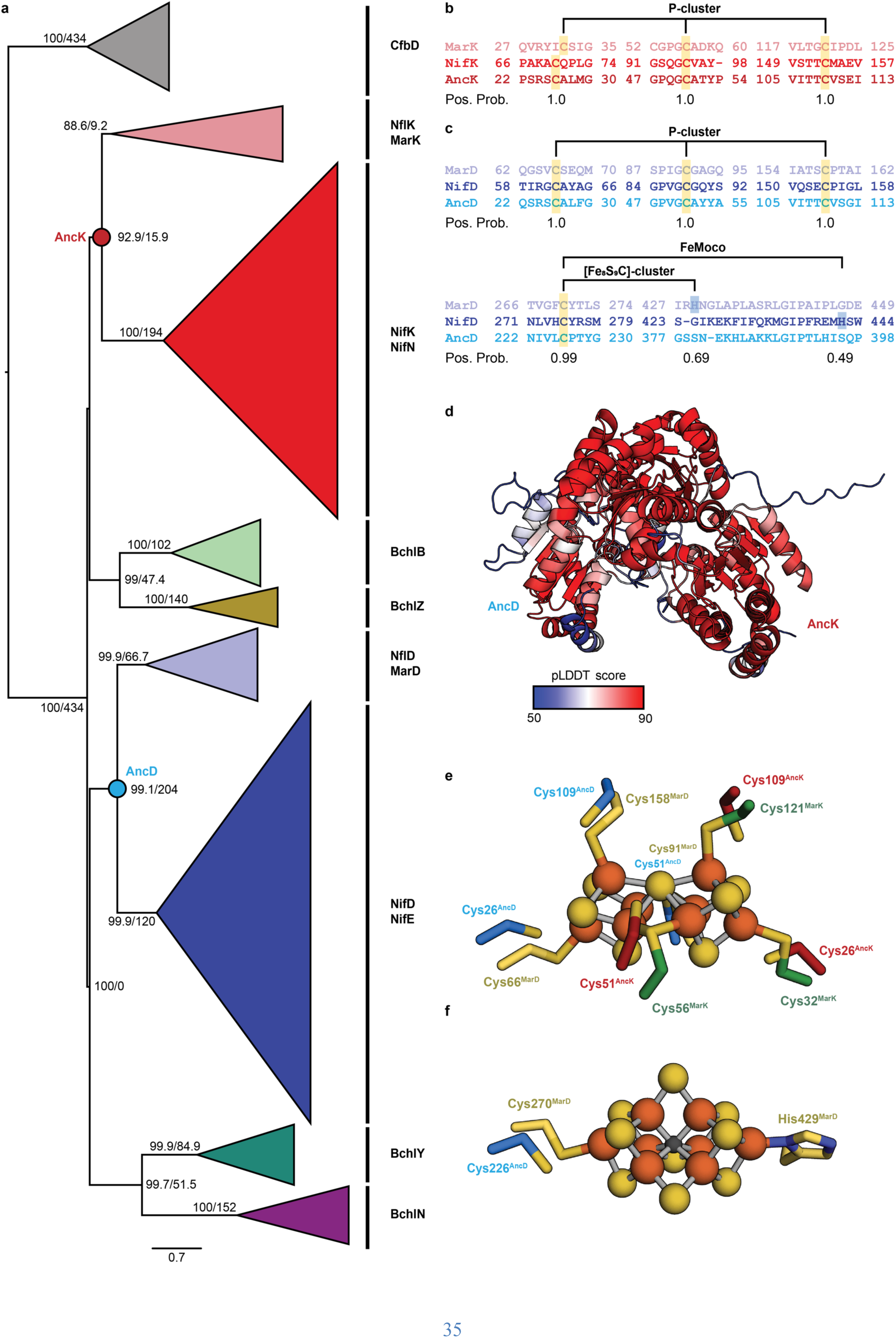
Phylogenetic relationship between the nitrogenase and the nitrogen fixation-like protein families. **a**, Maximum likelihood tree of Ni^2+^-sirohydrochlorin *a*,*c*-diamide reductive cyclase complex component D (CfbD), dark-operative protochlorophyllide *a* oxidoreductase (BchNB), chlorophyllide *a* oxidoreductase (BchYZ), nitrogenase-like proteins including the methylthio-alkane reductase (NflDK and MarDK) and Mo-dependent nitrogenase (NifDKEN) subunits. The tree was rooted with homomeric archaeal CfbD. The mirroring topology of nitrogenases and nitrogenase-like proteins implies an ancestral gene duplication giving rise to ancestral nitrogenase-like proteins and a second, more recent duplication for the emergence of bona fide Mo-nitrogenases (NifDKEN). The ancestral nodes resurrected correspond to the last common ancestor of MarD and NifD and the last common ancestor of NifK and MarK. Width of the clades is proportional to the number of sequences per clade. Supporting values represent the ultrafast bootstraps replicates by Iqtree2^63^ and the approximate likelihood-ratio (aLRT)^64^ statistical test, respectively. Scale bar represents the number of substitutions per site. **b,c**, Clustal Omega^68,69^ multiple sequence alignments (MSA) of the *Rhodospirillum rubrum* methylthio-alkane reductase MarK subunit to the *A. vinelandii* Mo-nitrogenase NifK subunit and the resurrected ancestral node sequence AncK (b) and *R. rubrum* MarD, *A. vinelandii* NifD and resurrected ancestral node sequence AncD (c), showing the binding sites for P-cluster, FeMoco and [Fe_8_S_9_C]-cluster. Pos. Prob., posterior probability. **d**, Structure of the AncDK dimer predicted by AlphaFold 2^65^ colored by predicted local distance difference (pLDDT) score. **e,f**, Close-up of the methylthio-alkane reductase P-cluster (e) and [Fe_8_S_9_C]-cluster (f) aligned to the predicted structure of the ancestor AncDK dimer. Amino acid residues coordinating the metalloclusters are shown in stick representation and colored in yellow, green, blue and red for MarD, MarK, AncD and AncK, respectively.

## Notes

### Competing Interest Statement

The authors have declared no competing interest.

