## Supplemental Information for "Methylthio-alkane reductases use nitrogenase metalloclusters for carbon-sulfur bond cleavage"

**The PDF file includes:**

Supplementary Discussion 1 to 4
Supplementary Tables 1 to 6
References

### Supplementary Discussion

Supplementary Discussion 1: The reductase component $MarH_2$ belongs to the family of P-loop NTPases

Analyzing the structure, we observed a canonical reductase component $MarH_2$ containing switch I and switch II regions, which facilitate conformational changes associated with nucleotide binding and hydrolysis [1] (Extended Data Fig. 4a). Each monomer binds a MgADP-$AlF_3$ moiety at the Walker A motif (GKGGIGKS) [2], indicating the ability of $MarH_2$ to bind and hydrolyze MgATP (Fig. 2b and Extended Data Fig. 4b). $MarH_2$ coordinates a single [$Fe_4S_4$]-cluster in its dimeric interface by $Cys97^{MarH}$ and $Cys133^{MarH}$ (Fig. 2c).

Supplementary Discussion 2: Comparison of P-cluster surroundings in Mo-nitrogenase and methylthioalkane reductase

The P-cluster of *A. vinelandii*'s Mo-nitrogenase undergoes structural rearrangements upon oxidation [3]. In the one-electron oxidized $P^{1+}$ state, the nitrogenase P-cluster symmetry is disrupted by the binding of the conserved $Ser188^{\mathit{Av}\mathrm{NifK}}$ to Fe6 [4], while an additional coordination of Fe5 by the backbone amide of $Cys88^{\mathit{Av}\mathrm{NifD}}$ occurs at the two-electron oxidized $P^{2+}$ state [5]. In the Mo-nitrogenase of *Gluconacetobacter diazotrophicus*, the serine is substituted by an unusual coordination between $Tyr98^{\mathit{Gd}\mathrm{NifK}}$ and Fe8 [6]. While MarK lacks structural analogs of both $Ser188^{\mathit{Av}\mathrm{NifK}}$ and $Tyr98^{\mathit{Gd}\mathrm{NifK}}$, $Cys52^{MarK}$ or $Gln60^{MarK}$ might coordinate Fe8, akin to $Tyr98^{\mathit{Gd}\mathrm{NifK}}$ (Extended Data Fig. 6b). Alternatively, P-cluster coordination upon oxidation could involve Fe4 and the residues $Ser157^{MarD}$ or $Glu68^{MarD}$. However, in the current structure all these potential ligands are at relatively long distances of 4.8-5.7 Å to the respective Fe atoms. Our findings strongly suggest that the P-cluster of methylthio-alkane reductase is in the reduced state ($P^N$), however, future studies on different redox states will be necessary.

Supplementary Discussion 3: EPR features of $MarH_2$

The temperature dependency of the two absorption-shaped peaks at $g$=5.79 and $g$=5.03 of the $[Fe_4S_4]^{1+}$ cluster $S$=3/2 state define the $S$=3/2 spin Hamiltonian parameters as $|E/D|$=0.20 and D≈ 2 $cm^{-1}$ in the absence of ATP (Extended Data Fig. 8d-g, and Extended Data Table 2). Structural changes induced by ATP decrease the $S$=1/2 content and the amplitude of the zero-field splitting parameter D of the $S$=3/2 species in the $MarH_2$ homodimer, in agreement with its function as ATP-dependent reductase.

Supplementary Discussion 4: Differences between EPR features of $Mar(DK)_2$ and $Nif(DK)_2$

We employed the catalytic component of the Mo-nitrogenase $Nif(DK)_2$ from *R. capsulatus* as EPR spectroscopic benchmark (Extended Data Fig. 9 and Extended Data Table 3). Though $S$=3/2 EPR signals of FeMoco in the DT-reduced state, a mixture of two very intense $S$=1/2 signals of the $P^{1+}$-cluster and the $g$=15.7 signal of the $P^{2+}$-cluster, were easily detected for $Nif(DK)_2$, no such signals were found for $Mar(DK)_2$ in various redox states. Although both methylthio-alkane reductase and nitrogenase contain a P-cluster, their electron transport mechanisms may differ since the characteristic $P^{2+}$ integer spin EPR signal [7] could not be detected. These differences likely originate from dissimilarities in the protein environments. The methylthio-alkane reductase lacks the conserved $Ser188^{\mathit{Av}\mathrm{NifK}}$ of nitrogenases, which is involved in P-cluster ligation upon oxidation.

**Supplementary Table 1: Information of cryo-EM data collection and model refinement**

| | **MarD$K_2H_2$ (EMD-50553), (PDB: 9FMG)** |
|---|---|
| **Data collection and processing** | |
| Magnification | 105,000 |
| Voltage (keV) | 300 |
| Electron exposure ($e^-/Å^2$) | 50 |
| Defocus range (μm) | −1.0 to −2.5 |
| Pixel size (Å) | 0.837 |
| Symmetry imposed | C1 |
| Initial particle images (no.) | 10,625,194 |
| Final particle images (no.) | 116,370 |
| Map resolution (Å) | 2.75 / 4.10 (masked / unmasked) |
| FSC threshold | 0.143 |
| Map resolution range (Å) | 1.8–12.4 |
| Map sharpening B factor ($Å^2$) | -71.5 |
| **Refinement** | |
| Initial model used | AlphaFold 2 |
| Model resolution (Å) | 2.71 / 2.74 (masked / unmasked) |
| FSC threshold | 0.143 |
| $CC_{Map}$ | 0.76 |
| Model composition | |
| Non-hydrogen atoms | 13,763 |
| Protein residues | 1,768 |
| Ligands | 2 × ADP, 2 x $AlF_3$, 2 × $Mg^{2+}$, 1 × [$Fe_8S_9C$], 1 × [$Fe_8S_7$], 1 × [$Fe_4S_4$] |
| B factors ($Å^2$) | |
| Protein (min / max / mean) | 15.55 / 108.68 / 42.94 |
| Ligand (min / max / mean) | 13.40 / 63.17 / 38.37 |
| R.m.s. deviations | |
| Bond lengths (Å) | 0.28 |
| Bond angles (°) | 0.51 |
| **Validation** | |
| MolProbity score | 1.71 |
| Clashscore | 6.00 |
| CaBLAM outliers | 2.53 |
| Poor rotamers (%) | 0.28 |
| C-beta deviations | 0.00 |
| Ramachandran plot | |
| Favored (%) | 94.47 |
| Allowed (%) | 5.53 |
| Disallowed (%) | 0.00 |

**Supplementary Table 2: Parameters for the simulation of *S*=3/2 EPR spectra of $MarH_2$**

| EPR species | T (K) | *g*-value | Amplitude | FWHM* | *g*-value | Amplitude | FWHM* | *g*-value | Amplitude† | FWHM*† |
|---|---|---|---|---|---|---|---|---|---|---|
| $MarH_2$ without ATP | 4.0 | 5.79 | 0.50 | 7.0 | 5.03 | 0.45 | 32 | 4.29 | 4.5 | 5.5 |
| | 10 | 5.79 | 0.18 | 7.0 | 5.03 | 0.42 | 32 | 4.29 | 5.0 | 5.5 |
| | 20 | 5.79 | 0.05 | 7.0 | 5.03 | 0.21 | 32 | 4.29 | 3.0 | 5.5 |
| $MarH_2$ with ATP | 4.0 | 5.76 ‡ | nd § | 10 | 5.06 | 1.07 | 28 | 4.28 | 7.0 | 5.5 |
| | 10 | 5.76 ‡ | nd § | 10 | 5.06 | 0.62 | 28 | 4.28 | 6.5 | 5.5 |
| | 20 | 5.76 ‡ | nd § | 10 | 5.06 | 0.23 | 28 | 4.28 | 4.0 | 5.5 |

* Full width at half maximum (FWHM) of the simulation with a Gaussian line shape.

† Amplitude and FWHM refer to the simulation before calculation of its derivative.

‡ Calculated *g*-value from the rhombogram using *g*=5.06 (see Extended Data Fig. 8).

§ Not detectable, possibly too broad.

**Supplementary Table 3: Parameters for the simulation with GeeStrain5 of *S*=1/2 EPR spectra**

| EPR species | $g_{av}$ | $g_x$ | $g_y$ | $g_z$ | $W_{xx}$ | $W_{yy}$ | $W_{zz}$ | $w_{xy}$ | $w_{xz}$ | $w_{yz}$ |
|---|---|---|---|---|---|---|---|---|---|---|
| Low potential* | 1.959 | 1.880 | 1.936 | 2.060 | 0.030 | 0.010 | 0.030 | 0.007 | 0.025 | 0 |
| L-cluster-like† | 1.908 | 1.830 | 1.926 | 1.967 | 0.095 | 0.041 | 0.022 | 0 | 0 | 0.006 |
| $P^{1+}$ major‡ | 1.955 | 1.841 | 1.962 | 2.062 | 0.008 | 0.006 | 0.007 | 0 | 0 | 0 |
| $P^{1+}$ minor‡ | 1.933 | 1.885 | 1.907 | 2.006 | 0.006 | 0.010 | 0.005 | 0 | 0 | 0 |

* For reduced $Mar(DK)_2$ in Fig. 2g (bottom trace).

† For oxidized $Mar(DK)_2$ in Fig. 2g (trace below $Nif(EN)_2$).

‡ For 0.5 mM IDS oxidized $Nif(DK)_2$ of *R. capsulatus* in Extended Data Fig. 9c. Ratio major to minor species: 1 to 0.20 (in GeeStrain5, corresponding to integrated intensity).

**Supplementary Table 4: Strains used in this study**

| Strain | Genotype | Reference |
|---|---|---|
| *Rhodospirillum rubrum* ATCC 11170 / S1 | Wildtype | Ref. [8] |
| *Rhodobacter capsulatus* B10S | Wildtype | Ref. [9] |
| *Rhodobacter capsulatus* MM0246 | *ΔnifD::SpR ΔanfHDGK::gmR ΔdraTG ΔmodABC ΔgtaI* | Ref. [10] |
| *Rhodobacter capsulatus* MM0422 | *ΔnifD::SpR ΔanfHDGK::gmR ΔdraTG ΔmodABC ΔgtaI* / pMM0181 | This study |
| *Rhodobacter capsulatus* MM0468 | *ΔnifHDK ΔanfHDGK::GmR ΔdraTG ΔgtaI* | Ref. [11] |
| *Rhodobacter capsulatus* MM0480 | *ΔnifHDK ΔanfHDGK::GmR ΔdraTG ΔgtaI* / pMM0207 | Ref. [11] |
| *Escherichia coli* DH5α | $F^-$ Φ80*lac*ZΔM15 Δ(*lac*ZYA-*arg*F) U169 *rec*A1 *end*A1 *hsd*R17($r_k^-$, $m_k^+$) *pho*A *sup*E44 *thi*-1 *gyr*A96 *rel*A1 $\lambda^-$ | Thermo Fisher Scientific, (Waltham, Massachusetts, USA) catalogue #18265017 |
| *Escherichia coli* ST18 | *pro thi hsdR*$^+$ $Tp^r$ $Sm^r$; chromosome::RP4-2 Tc::Mu-Kan::Tn7/lpir *ΔhemA* | Ref. [12] |

**Supplementary Table 5: Primers used in this study**

| Primer | Sequence | Purpose |
|---|---|---|
| oMM0494 | CACCACAGGTCTCGTATGACGGTTCCTGCTTATCCTTCC | Construction of pMM0165 |
| oMM0495 | CACCACAGGTCTCGAGCGTCAAGCGCTTGCGCTGA | Construction of pMM0165 |
| oMM0649 | CGCTCTTGGACTCCTG | Construction of pMM0170 and pMM0181 |
| oMM0650 | GTTAAACAAAATTATTTCTAGACGGC | Construction of pMM0170 and pMM0181 |
| oMM0651 | GGCCGTCTAGAAATAATTTTG | Construction of pMM0181 |
| oMM0652 | CTGGATCTATCAACAGGAGTC | Construction of pMM0170 and pMM0181 |
| oMM0662 | ATGGTGATGATGGTGGTGCATGGTTCTCTCCGTC | Construction of pMM0181 |
| oMM0663 | CACCACCATCATCACCATGCCAAAAGTCCCAAAC | Construction of pMM0181 |
| oMM0668 | CACCTGGCCGTCTAGAAATAATTTTG | Construction of pMM0170 |
| oMM0669 | TTTTTCGAACTGCGGGTGGCTCCACTCTGCGGGACGGCG | Construction of pMM0170 |
| oMM0670 | TGGAGCCACCCGCAGTTCGAAAAATGAGCCCCGTCATGC | Construction of pMM0170 |

**Supplementary Table 6: Plasmids used in this study**

| Plasmid | Code | Description | Reference |
|---|---|---|---|
| pOGG024 | pMM0114 | Broad-host range and medium copy number, pBBR1 with *oriT*, *lacZα* cassette for golden gate cloning (BsaI), $Gen^R$ | Ref. [13] |
| pOGG024-*kanR* | pMM0119 | Broad-host range and medium copy number, pBBR1 with *oriT*, *lacZα* cassette for golden gate cloning (BsaI), $Kan^R$ | Ref. [10] |
| pOGG024-*kanR* | pMM0129 | Broad-host range and medium copy number, pBBR1 with *oriT*, *lacZα* cassette for golden gate cloning (BsaI), *anfH* promoter, $Kan^R$ | Ref. [14] |
| pOGG024-*kanR marBHDK* | pMM0165 | Broad-host range and medium copy number, pBBR1 with *oriT*, *marBHDK* cloned into BsaI site, *anfH* promoter, $Kan^R$ | This study |
| pOGG024-*kanR marBHDK*-Strep | pMM0170 | Broad-host range and medium copy number, pBBR1 with *oriT*, *marBHDK* cloned into BsaI site with C-terminal Strep tag II on *marD*, *anfH* promoter, $Kan^R$ | This study |
| pOGG024-*kanR marBHDK*-Strep/His | pMM0181 | Broad-host range and medium copy number, pBBR1 with *oriT*, *marBHDK* cloned into BsaI site with N-terminal hexahistidine tag on *marH* and C-terminal Strep tag II on *marD*, *anfH* promoter, $Kan^R$ | This study |
| pOGG024-*kanR nifHDK*-Strep/His | pMM0207 | Broad-host range and medium copy number, pBBR1 with *oriT*, *nifHDK* cloned into BsaI site with N-terminal hexahistidine tag on *nifH* and C-terminal Strep tag II on *nifD*, *nifH* promoter, $Kan^R$ | Ref. [11] |
